# A Metric Space of Ranked Tree Shapes and Ranked Genealogies

**DOI:** 10.1101/2019.12.23.887125

**Authors:** Jaehee Kim, Noah A. Rosenberg, Julia A. Palacios

**Affiliations:** Department of Biology, Stanford University, Stanford, CA 94305, USA; Department of Statistics, Stanford University, Stanford, CA 94305, USA; Department of Biomedical Data Science, Stanford Medicine, Stanford, CA 94305, USA

## Abstract

Genealogical tree modeling is essential for estimating evolutionary parameters in population genetics and phylogenetics. Recent mathematical results concerning ranked genealogies without leaf labels enable new opportunities in the analysis of evolutionary trees. In particular, comparisons between ranked genealogies facilitate the study of evolutionary processes for organisms sampled in multiple time periods. We propose a metric space on ranked genealogies for lineages sampled from both isochronous and time-stamped heterochronous sampling. Our new tree metrics make it possible to conduct statistical analyses of ranked tree shapes and timed ranked tree shapes, or ranked genealogies. Such analyses allow us to assess differences in tree distributions, quantify estimation uncertainty, and summarize tree distributions. We show the utility of our metrics via simulations and an application in infectious diseases.

## Introduction

Gene genealogies are rooted binary trees that describe the ancestral relationships of a sample of molecular sequences at a locus. The properties of these genealogies are influenced by the nature of the evolutionary forces experienced by the sample’s ancestry. Hence, assessing differences between gene genealogies can provide information about differences in these forces. In this manuscript, we propose a distance on genealogies that enables biologically meaningful comparisons between genealogies of non-overlapping samples. Our proposed distance, in its more general form, is a distance on *ranked genealogies*.

Ranked genealogies are rooted binary and unlabeled trees with branch lengths and ordered internal nodes (Figure 1(A)). Ranked genealogies are also known as Tajima’s genealogies (Sainudiin et al., 2015; Palacios et al., 2019) as they were examined by Tajima (1983). Ranked genealogies are coarser than labeled topologies with branch lengths specified but finer than unranked unlabeled genealogies in which neither branch order nor taxon labels are specified (Figure 1). Recently, there has been increasing interest in modeling ranked genealogies for studying evolutionary dynamics (Ford et al., 2009; Lambert and Stadler, 2013; Sainudiin et al., 2015). A new method for inferring evolutionary parameters from molecular data is based on the Tajima coalescent of ranked genealogies (Palacios et al., 2019): modeling of ranked unlabeled genealogies, as opposed to ranked labeled genealogies (Kingman coalescent), reduces the dimensionality of the inference problem. In studying macroevolution, Maliet et al. (2018) proposed a model on ranked unlabeled tree topologies to investigate factors influencing nonrandom extinction and the loss of phylogenetic diversity.

**Figure 1:**
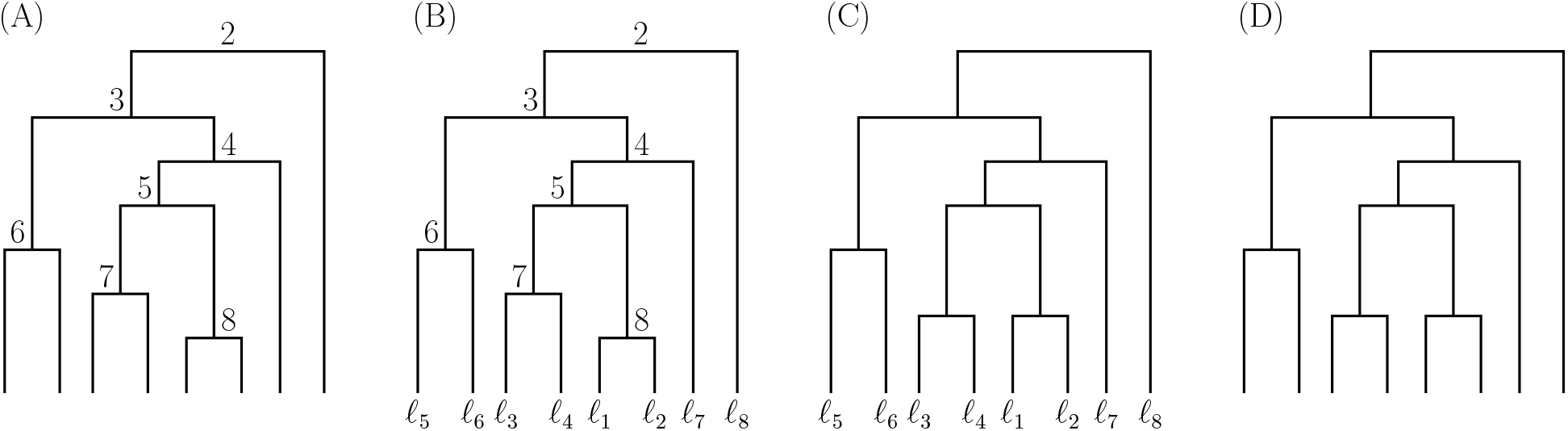
Types of tree topology. (A) Ranked tree shape (*T*^*R*^). (B) Labeled ranked tree shape (*T*^*LR*^). (C) Labeled unranked tree shape (*T*^*L*^). (D) Tree shape (*T*^*S*^). Genealogies corresponding to each topology include branch lengths.

Many metrics on labeled binary trees—such as the Robinson-Foulds (RF) metric (Robinson and Foulds, 1981), the Billera-Holmes-Vogtmann (BHV) metric (Billera et al., 2001), and the Kendall-Colijn metric (Kendall and Colijn, 2016)—have been proposed. These metrics make possible comparisons between labeled genealogies. They have been used for summarizing posterior distributions and bootstrap distributions of trees on the same set of taxa (Chakerian and Holmes, 2012; Brown and Owen, 2019), for comparing estimated genealogies of the same organisms with different procedures, and for quantifying uncertainty (Willis and Bell, 2018). A metric on the lower resolution space of tree shapes—unlabeled unranked binary trees without branch lengths—has recently been proposed by Kendall and Colijn (2016). To our knowledge, no other metrics on ranked unlabeled genealogies have been proposed to date.

A tree metric on the space of ranked unlabeled genealogies facilitates evaluations of the quality of an estimation procedure, by enabling measurements of the distance between an estimated ranked genealogy and the true ranked genealogy. It can assist in comparing different estimators from different procedures, and in comparing estimated ranked genealogies of different organisms at different geographical locations and different time periods. Moreover, a useful tree metric not only provides a quantitative measure for ranked genealogy comparison, it can also discriminate between the key aspects of different evolutionary processes (Holmes, 2003). We show that our metrics separate samples of genealogies originating from different distributions of ranked tree topologies and ranked genealogies. Our distances enable the computation of summary statistics, such as the mean and the variance, from samples of ranked genealogies. When ranked genealogy samples are obtained from posterior distributions of genealogies, such as those obtained from BEAST (Suchard et al., 2018), our tree metrics enable the construction of credible sets.

We first define a metric on ranked unlabeled tree shapes. Our metric relies on an integer-valued triangular matrix representation of ranked tree shapes. This matrix representation allows us to use metrics on matrices, such as the Frobenius norm, to define metrics on ranked tree shapes. The choice of the Frobenius norm produces computational benefits, as computations of the metrics are quadratic in the number of leaves. Our metrics on ranked unlabeled tree shapes retain more information than metrics based on unlabeled unranked tree shapes alone.

We expand our metric definition to ranked genealogies including branch lengths by weighting the matrix representation of ranked tree shapes by branch lengths. We first define a distance on ranked unlabeled isochronous trees, with all samples obtained at the same point in time. We then later define a metric space on heterochronous ranked tree shapes and heterochronous genealogies, in which samples are time-stamped. While modeling isochronous genealogies is the standard practice for slowly evolving organisms, such as humans and other types of animals, modeling heterochronous genealogies is the standard approach for rapidly evolving organisms, such as viruses and other pathogens. Heterochronous genealogies are also increasingly relevant in the study of ancient DNA samples.

We analyze different properties of our proposed metrics and compare them to other tree-valued metrics— such as BHV (Billera et al., 2001) and Kendall-Colijn (Kendall and Colijn, 2016)—by projecting these other metrics into the space of ranked tree shapes and ranked genealogies. We then demonstrate the performance of our metrics on simulations from various tree topology distributions and demographic scenarios, and we use them to compare posterior samples of genealogies of human influenza A virus from different geographic regions.

## Preliminaries

### Definitions of tree topologies and genealogies

All the trees we consider are rooted and binary. We first assume that trees are isochronously sampled, that is, all tips of the trees start at the same time. A *ranked tree shape* with *n* leaves is a rooted unlabeled binary tree with an increasing ordering of the *n* − 1 interior nodes, starting at the root with label 2 (Figure 1(A)). We use the symbol *T*^*R*^ to denote a ranked tree shape. A *ranked genealogy* is a ranked tree shape equipped with branch lengths. We use the symbol **g**^*R*^ to denote a ranked genealogy. Although we mainly focus on ranked tree shapes and ranked genealogies, we will compare our metrics to metrics defined on other rooted tree spaces. A *labeled ranked tree shape* (Figure 1(B)) and a *labeled ranked genealogy* are the corresponding labeled counterparts of unlabeled ranked trees, and they are denoted by *T*^*LR*^ and **g**^*LR*^ respectively. A *labeled unranked tree shape* (*T*^*L*^) (Figure 1(C)) and *labeled unranked genealogy* (**g**^*L*^) are a rooted labeled tree shape and a rooted labeled genealogy without ranking of internal nodes, respectively. A *tree shape* (Figure 1(D)) is an unlabeled unranked rooted bifurcating tree denoted by *T*^*S*^ and the corresponding *genealogy* is denoted by **g**^*S*^.

### Probability models on tree topologies and genealogies

Generative probability models on *isochronous* binary tree topologies with *n* leaves are Markov jump processes that either start at the root with two branches and proceed (forward in time) by choosing one of the extant branches to bifurcate at random until the desired number of leaves is obtained, or start at the bottom with *n* leaves and proceed (backward in time) by choosing two extant lineages to merge into a single lineage until there is a single lineage. The first class corresponds to branching processes (Aldous, 1996; Ford, 2005; Steel, 2016) and the second class corresponds to coalescent jump processes (Kingman, 1982; Wakeley, 2009). The corresponding *heterochronous* Markov jump forward and backward in time processes are the birth-death process and the heterochronous coalescent process, respectively. In the heterochornous coalescent, sampling times (death of lineages) are usually assumed fixed (Felsenstein and Rodrigo, 1999). In each case, the distribution of branching times or coalescent times is independent of the tree topology and depends only on the number of extant branches.

In this manuscript, we simulate isochronous ranked tree shapes from two different models: the one-parameter beta-splitting model (Aldous, 1996) and a two-parameter model which we term the alpha-beta splitting model (Maliet et al., 2018). To generate isochronous ranked genealogies and heterochronous ranked tree shapes and ranked genealogies, we use the Tajima coalescent formulation of the coalescent (Sainudiin et al., 2015; Palacios et al., 2019). In the following, we briefly describe these models. Extensive accounts of probability models of evolutionary trees can be found in (Mooers and Heard, 1997; Ford, 2005; Blum and François, 2006; Steel, 2016; Sainudiin and Véber, 2016).

### Beta-splitting model on labeled tree shapes

We first consider the single-parameter *beta-splitting model* on labeled tree shapes (Aldous, 1996). For a parent clade of size *n*, the model chooses its child clade size to be *i* on the left branch and *n − i* on the right branch with probability

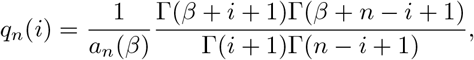

where *a_n_*(*β*) is a normalizing constant and *i* ∈ {1, 2, …, *n* − 1}. The splitting is repeated recursively in each branch independently until the tree is fully resolved. The parameter *β ∈* [*−*2, *∞*) controls the degree of balance of the generated trees. With *β* = *−*2, the model generates the perfect unbalanced tree (caterpillar tree) with probability one, and with *β* = *∞*, the model generates the perfect balanced tree with probability one. In particular, we consider the following special cases: the Yule model (Yule, 1925; Harding, 1971) (*β* = 0), the proportional-to-distinguishable-arrangements (PDA) model (Blum and François, 2006) (*β* = *−*1.5), and the Aldous’ branching (AB) model (Aldous, 2001) (*β* = *−*1). Additionally, we include *β* = *−*1.9 and *β* = 100, to which we refer as “unbalanced” and “balanced”, respectively. We note that Ford’s alpha-model Ford (2005) is another single parameter family of models on the same class of trees as the beta-splitting model and it is not considered in this manuscript.

### Alpha-beta splitting model and beta-splitting model on ranked tree shapes

The *alpha-beta model* of Maliet et al. (2018) generates labeled ranked tree shapes according to a size-biased distribution with a stick-breaking construction. The algorithm first generates *n* independent draws *u*_1_, …, *u_*n*_* from a Unif(0, 1) distribution. The *u_i_*’s correspond to the *n* leaves of the tree. The first partition of the *n* leaves (at the root) is determined by drawing a random number *R*_1_ *∼* Beta(*β* + 1, *β* + 1). All the 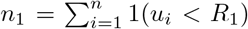 are placed on the left side of the tree and the rest on the other side. Then, if *n*_1_ > 1, the left branch is chosen to bifurcate with probability proportional to 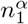. The algorithm continues generating beta-distributed values to bi-partition the leaves and chooses the order (ranking) proportional to their number of descendants until the interval (0, 1) is partitioned into *n* intervals, each corresponding to an *u_i_* number. The pseudocode is in Section S1. Labels are then removed to generate a ranked tree shape. The *β ∈* [*−*2, ∞) parameter determines the balance of the tree as in the beta-splitting model, and the *α ∈* (*−∞, ∞*) parameter regulates the relationship between subtree family size (number of descendants) and closeness to the root. More specifically, when *α* < 0, subtrees with small family sizes are closer to the root and when *α* > 0, subtrees with small family sizes are closer to the tips. When *α* = 1, the alpha-beta model becomes a *Beta-splitting model on ranked tree shapes*.

### Tajima coalescent on ranked genealogies

The Tajima coalescent is a model on ranked genealogies (Figure 1(A)). It is a Markov lumping of Kingman’s *n*-coalescent on labeled and ranked genealogies (Sainudiin et al., 2015; Palacios et al., 2019). The Tajima coalescent is a pure death process that starts with *n* unlabeled leaves at time *t_n_*_+1_ = 0 and proceeds backward in time, merging pairs of branches to create interior nodes labeled by their order of appearance. The merging of two branches is a coalescent event. In the Tajima coalescent, the distribution of coalescence times is the same as in the Kingman coalescent and the probability of a topology, ranked tree shape, is given by

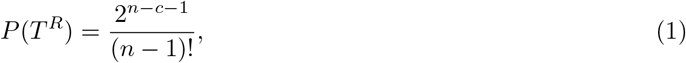

where *n* is the number of leaves, and *c* is the number of cherries—the number of pairs of leaves that subtend from a shared interior node. The Tajima coalescent on ranked tree shapes without times corresponds to the beta-splitting model on ranked tree shapes with *β* = 0, also called the Yule model. A full description of the Tajima coalescent process can be found in Cappello and Palacios (2019).

### Distributions on branching or coalescent times

As mentioned before, in this manuscript we consider tree models of neutral evolution in which branching or coalescent waiting times depend on the number of extant branches but are independent of the particular topology configuration or clade composition. A popular model in phylogenetics is to assume that, when there are *k* branches, the branching waiting time forward in time from root to tips is exponentially distributed with rate *kλ* with *λ* > 0 (Mooers and Heard, 1997). We will refer to this model as *λ-branching*. In the standard coalescent model, when there are *k* lineages, the coalescent time *t_k_* backward in time from tips (*k*) to root (see Figure 3(A)) is exponentially distributed with rate 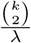, where *λ* is referred to as the effective population size, which is assumed to be constant over time. We will refer to this model as the *λ-coalescent*. In the coalescent with variable population size (Slatkin and Hudson, 1991), the first coalescent time when there are *n* lineages has marginal density

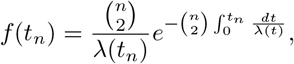

and the conditional density of *t_k_* when there are *k* branches is

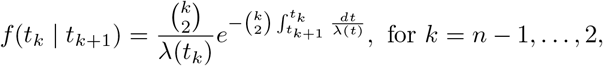

where *λ*(*t*) > 0 is the effective population size at time *t*. We will refer to the variable population size distribution on coalescent times as the *λ*(*t*)*-coalescent*.

**Figure 2:**
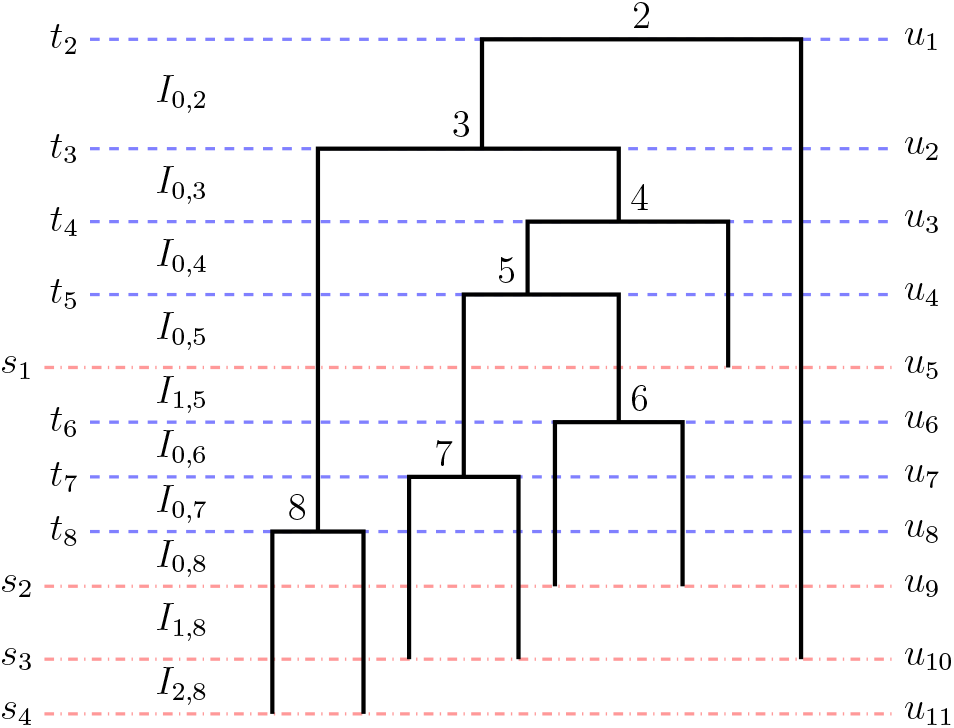
Heterochronous genealogy. Example of a ranked genealogy with heterochronous sampling. *s*_4_, …, *s*_1_ and *t*_8_, …, *t*_2_ indicate sampling times (red dotted lines) and coalescent times (blue dotted lines), respectively. *u*_11_, …, *u*_1_ are the ordered times of change points where the number of lineages changes due to either a sampling event or a coalescent event. The time increases backwards in time starting with *u*_11_ = 0 as the present time. *I*_0,*k*_ is the interval that ends with a coalescent event at *t_k_*. *I*_*i,k*_ (*i* > 0) represents the interval that ends with a sampling time within the interval (*t_k_*_+1_, *t*_*k*_). For *k* = *n*, we adopt the convention *t_n_*_+1_ = 0.

**Figure 3:**
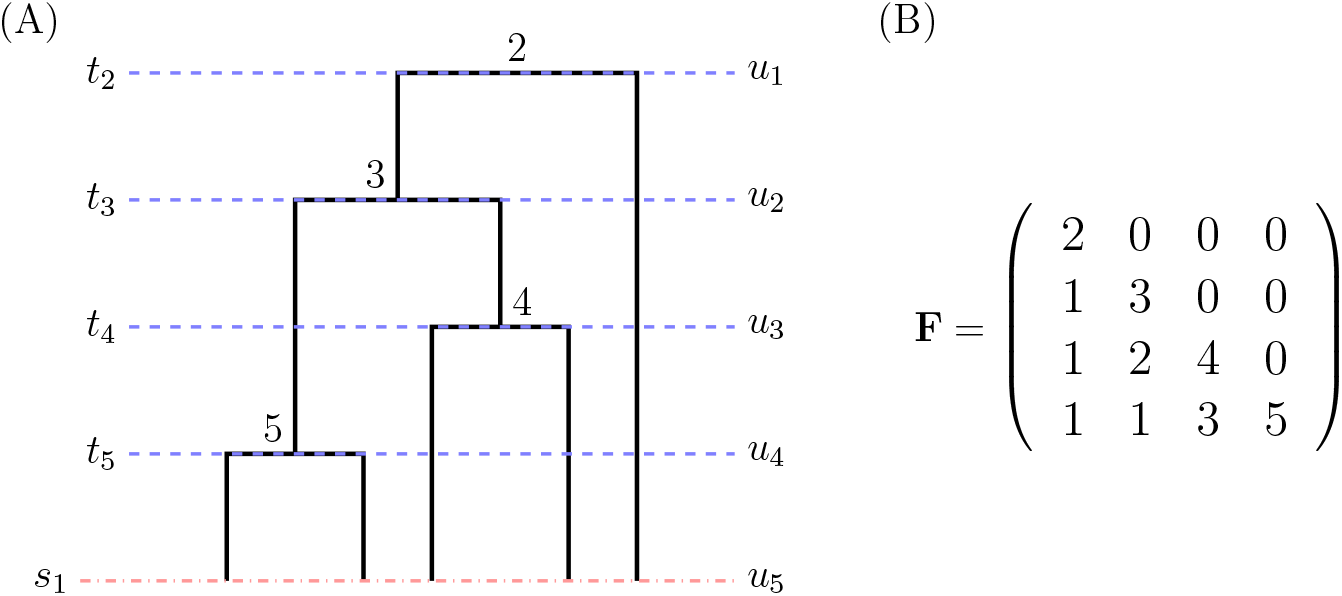
Unique encoding of isochronous ranked tree shapes and F-matrices. (A) Example of a ranked genealogy with isochronous sampling. (B) The corresponding **F**-matrix that encodes the ranked tree shape information of the tree in (A).

### Coalescent distribution on heterochronous tree topologies and genealogies

The ranked tree shape and ranked genealogy of time-stamped samples are termed *heterochronous ranked tree shapes* and *heterochronous genealogies*, respectively (Figure 2). Here, we assume that samples are collected at times *s_m_, s_m−_*_1_, …, *s*_1_, with *s_m_* = *t_n_*_+1_ = 0 (the present), and *s*_*j*_ < *s*_*j*−1_ for *j* = *m*, …, 2. At time *s_j_*, *n*_*j*_ lineages are sampled, and 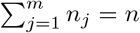.

The *λ(t)-heterochronous-coalescent* (Felsenstein and Rodrigo, 1999) describes the distribution of coalescent times conditioned on collecting *n_m_, n_m−_*_1_, …, *n*_1_ samples at times *s_m_, s_m−_*_1_, …, *s*_1_. As before, *t*_*n*+1_, *t*_*n*_, …, *t*_2_ denote the coalescent times, except that the subindex no longer indicates the number of lineages. Instead, the subindex indicates the rank order of the coalescent events going forward in time from the root at *t*_2_. Define (*u_n_*_+*m−*1_, *u*_*n*+*m−*2_, …, *u*_1_) as the vector of change points (coalescent or sampling times), with 0 = *u_n_*_+*m−*1_ < *u*_*n*+*m*−2_ < *· · ·* < *u*_1_ = *t*_2_. To state the density of coalescent times according to the *λ*(t)-heterochronous-coalescent, going backwards in time, we denote the intervals that end with a coalescent event at *t_k_* by *I*_0,*k*_, and the intervals that end with a sampling time within the interval (*t_k_*_+1_, *t*_*k*_) as *I*_*i,k*_, where *i* is an index tracking the sampling events in (*t*_*k*+1_, *t*_*k*_). More specifically,

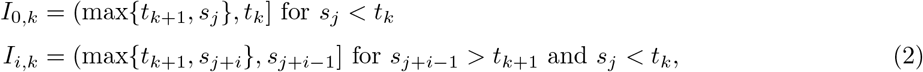

with *k* = 2, …, *n* and *i* ranges from 1 to the number of sampling events in (*t_k_*_+1_, *t*_*k*_). An example of the annotated time intervals using *I*_0,*k*_ and *I_i,k_* is shown in Figure 2.

The conditional density of *t_k−_*_1_ is the product of the conditional density of *t_k−_*_1_ *∈ I*_0,*k*_ and the probability of not having a coalescent event during the period of time spanned by intervals *I*_1,*k*_, …, I_*m,k*_. That is, for *k* = 2, …, *n*,

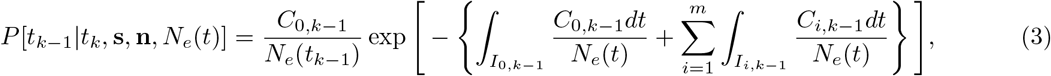

where 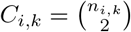.

### Tree metrics for comparative analysis

#### Metrics on labeled trees

A large number of tree metrics have been proposed for phylogenetics. Billera et al. (2001) introduced a metric space of trees for labeled genealogies now commonly known as BHV space. Owen and Provan (2011) and Chakerian and Holmes (2012) provided polynomial algorithms and implementations for calculating the geodesic distance metric proposed by Billera et al. (2001). Recently, Kendall and Colijn (2016) proposed a new metric on labeled unranked trees and labeled genealogies representing each tree as a convex combination of two vectors—one vector encoding number of edges from the root to the most recent common ancestor of every pair of tips in the tree, and the other vector encoding the corresponding path lengths. The most popular metric that can be computed in polynomial time is the symmetric difference of Robinson and Foulds (RF) on labeled unranked tree shapes (Robinson and Foulds, 1981) and the branch-length measure RFL on labeled genealogies (Robinson and Foulds, 1979). Other metrics on labeled tree shapes are based on the number of rearrangement steps needed to transform one tree into the other, including the nearest neighbor interchange distance (Li and Zhang, 1999) and the metrics based on pruning and regrafting or tree bisection and reconnection (Allen and Steel, 2001; Bordewich and Semple, 2005).

#### Metrics on unlabeled trees

A metric on unlabeled trees is desirable because it allows comparison between trees from different samples. However, not many such metrics are available. Colijn and Plazzotta (2018) proposed a metric on tree shapes which is the Euclidean norm of the difference between two integer vectors that uniquely describe the two trees. Poon et al. (2013) developed a kernel function that measures the similarity between two unlabeled genealogies by accounting for differences in branch lengths and matching number of descendants over all nodes for both trees. Lewitus and Morlon (2015) proposed the Jensen-Shannon distance between the spectral density profiles of the corresponding modified graph Laplacian of the unlabeled genealogies. The modified graph Laplacian of an unlabeled genealogy is constructed as the difference between its degree matrix—a diagonal matrix with *i*-th diagonal element being the sum of the branch lengths from node *i* to all the other nodes—and its distance matrix whose (*i, j*)-element is the branch length between nodes *i* and *j*.

To our knowledge, we are introducing the first metric for unlabeled ranked tree shapes and unlabeled ranked genealogies. In order to define our metric on ranked tree shapes, we need to introduce a unique encoding of a ranked tree shape as an integer-valued triangular matrix defined in the next section.

## New Approaches

### Unique encoding of ranked tree shapes and F-matrix

Let *T*^*R*^ be a ranked tree shape with *n* leaves sampled at time 0 = *u_n_*. Let (*u_n−_*_1_, …, *u*_1_) be the *n −* 1 coalescent times. Here, time increases into the past, *u_n−_*_1_ < *· · ·* < *u*_1_, and *u_n_* and *u*_1_ correspond to the most recent sampling time and the time to the most recent common ancestor (root), respectively. An **F**-matrix that encodes *T*^*R*^ is an (*n −* 1) *×* (*n −* 1) lower triangular matrix of integers with elements *F_i,j_* = 0 for all *i* < *j*, and for 1 *≤ j ≤ i*, *F_i,j_* is the number of extant lineages in (*u_j_*_+1_, *u_j_*) that do not bifurcate during the entire time interval (*u_i_*_+1_, *u_j_*).

In Figure 3(B), we show the corresponding **F**-matrix to the ranked tree shape depicted in Figure 3(A). In the interval (*u*_2_, *u*_1_), there are two lineages so *F*_1,1_ = 2. One of the two lineages extant at time (*u*_2_, *u*_1_) branches at time *u*_2_ while the other lineage does not branch throughout the entire time interval (*u*_5_, *u*_1_). This gives the first column of the **F**-matrix: *F*_2,1_ = *F*_3,1_ = *F*_4,1_ = 1. For the second column, we start with three lineages in the interval (*u*_3_, *u*_2_), *F*_2,2_ = 3. Of the three lineages extant at (*u*_3_, *u*_2_), one branches at *u*_3_ (*F*_3,2_ = 2), and one branches at *u*_4_ (*F*_4,2_ = 1). We construct the third column by starting from four lineages in (*u*_4_, *u*_3_), *F*_3,3_ = 4. One lineage branches at *u*_4_, and thus, *F*_4,3_ = 3. Finally, in the interval (*u*_5_, *u*_4_), there are five lineages, which gives *F*_4,4_ = 5.

If *T*^*R*^ is a heterochronous ranked tree shape with *n* leaves and *m* sampling events, the corresponding **F**-matrix representation is an (*n* + *m −* 2) *×* (*n* + *m −* 2) lower triangular matrix of integers, where *F_i,j_* for 1 *≤ j ≤ i* is the number of extant lineages in (*u_j_*_+1_, *u_j_*) that do not bifurcate or become extinct during the entire time interval (*u_i_*_+1_, *u_j_*) traversing forward in time. Here, (*u_n_*_+*m−*1_, *u_n_*_+*m−*2_, …, *u*_1_), such that 0 = *u_n_*_+*m−*1_ < *u*_*n*+*m*−2_ < *· · ·* < *u*_1_, are the *n* + *m −* 1 ordered change points of *T*^*R*^, at each of which the number of branches changes either by a coalescent event or by a sampling event. We show an example of a heterochronous ranked tree shape and its **F**-matrix encoding in Figure S2(C-D).

Although the branch lengths and the actual values of coalescent and sampling times are irrelevant for the specification of the ranked tree shape, we rely on the *u_i_* to identify the order and type of change points, coalescence or sampling, in *T*^*R*^ and **F**.

#### Theorem 1.

**(Unique Encoding of Ranked Tree Shapes).** The map by which ranked tree shapes with *n* samples and *m* sampling events are encoded as **F**-matrices of size (*n* + *m −* 2) *×* (*n* + *m −* 2) uniquely associates a ranked tree shape with an **F**-matrix.

In other words, given an **F**-matrix, if it encodes a ranked tree shape, it encodes exactly one ranked tree shape. The proof can be found in Section S2. In the next section, we will leverage the **F**-matrix representation of ranked tree shapes to define a distance between ranked tree shapes. From this point onward, we will assume a matrix **F** represents a ranked tree shape.

### Metric spaces on ranked tree shapes and ranked genealogies

We define two distance functions *d*_1_ and *d*_2_ on the space of ranked tree shapes with *n* leaves as follows. For a pair of ranked tree shapes 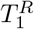 and 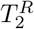 and their corresponding **F**-matrix representations **F**^(1)^ and **F**^(2)^ of size *r* × *r*,

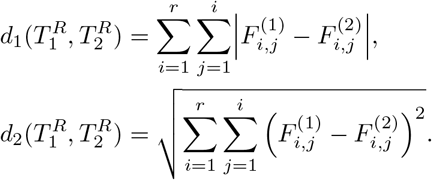

The two distances are metrics since they inherit properties of the *L*_1_-norm (Manhattan distance) and *L*_2_-norm (Frobenius norm). The definitions are valid for both isochronous and heterochronous ranked tree shapes as both types of trees have unique matrix encodings.

Figure 4 shows all isochronous ranked tree shapes of *n* = 5 leaves and their pairwise *d*_1_ distances. The *d*_1_ metric shows the following desirable features: the largest distances occur between trees with different tree shapes and between the most balanced and unbalanced trees. There are 3 different values of *d*_1_ among the 10 possible pairs of trees: 1, 2, and 3. The *d*_2_ distance exhibits the same features. The three different values of *d*_2_ pairwise distances are 1, 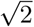 and 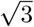, and the relative comparisons remain the same as for *d*_1_. In the following definition, we include branch lengths to define distances between ranked genealogies.

**Figure 4:**
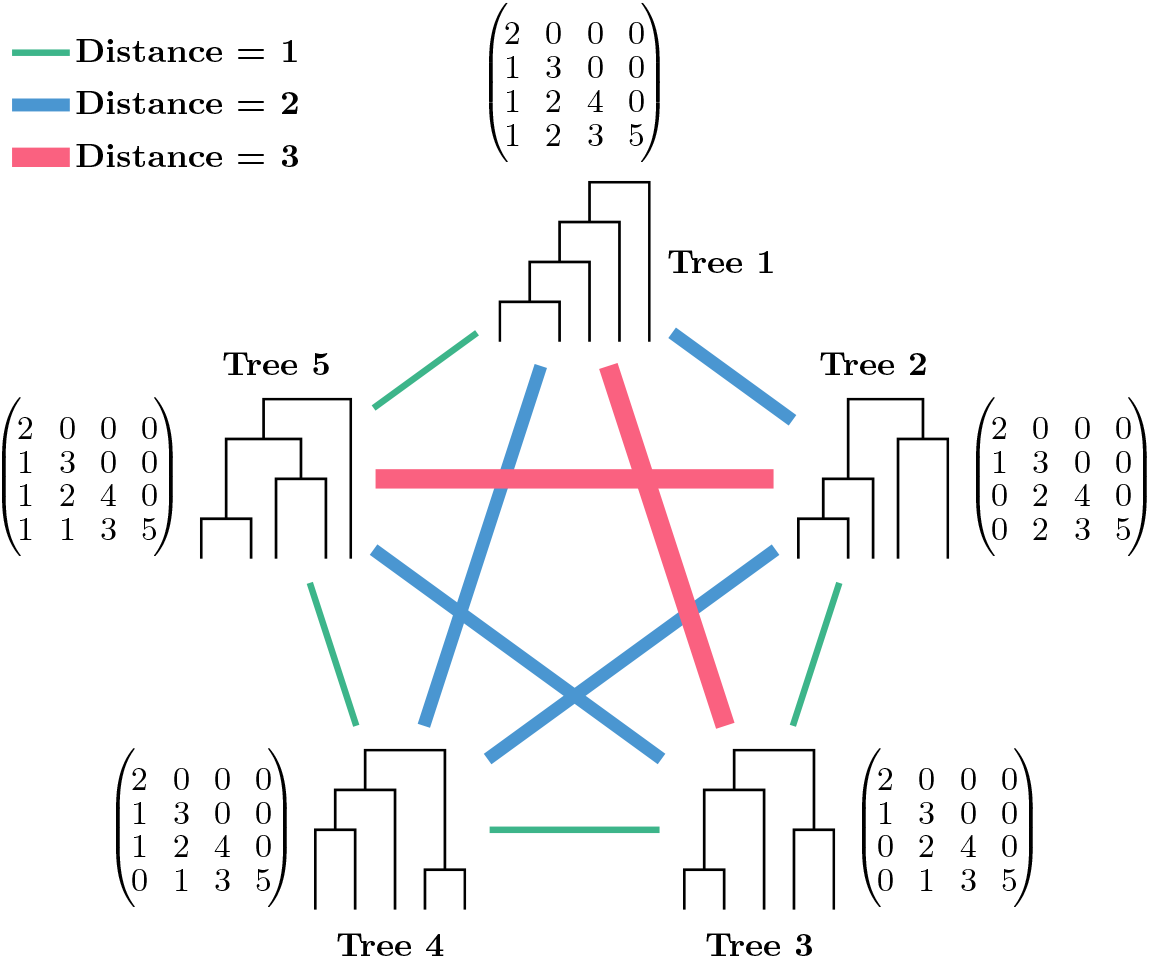
*d*_1_ distance between all pairs of ranked tree shapes of *n* = 5 leaves. Rankings of internal nodes are removed for ease of visualization. There are three different distance values among the 10 pairs of distances. The maximum distance of 3 is between trees 2 and 5 (both trees have different shapes) and between trees 1 and 3 (most unbalanced tree and most balanced tree). Trees 2, 3, and 4 share the same shape but with different internal node rankings. Among those three trees, Trees 2 and 4 differ by two ranking moves, whereas the other tree pairs differ by one ranking move. Indeed, trees 2 and 4 have distance 2, and the other two pairs have distance 1 under *d*_1_ metric.

We next define two weighted distance functions 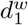 and 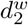 on the space of ranked genealogies with *n* samples and *m* sampling events. We first define the weight matrix **W** of a ranked genealogy **g**^*R*^ as a lower triangular matrix of size (*n* + *m −* 2) *×* (*n* + *m −* 2) with entries *W_i,j_* = *u_j_ − u_i_*_+1_ for *j ≤ i* and *W_i,j_* = 0 otherwise. Here, (*u_n_*_+*m−*1_, *u_n_*_+*m−*2_, …, *u*_1_), such that 0 = *u_n_*_+*m−*1_ < *u_n_*_+*m−*2_ < *· · ·* < *u*_1_, is the vector of ordered *n* − 1 coalescent times *t_k_* and *m* sampling times *s*_*ℓ*_ with time increasing into the past. For a pair of ranked genealogies 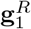 and 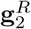, and their corresponding **F**-matrix representations **F**^(1)^ and **F**^(2)^,

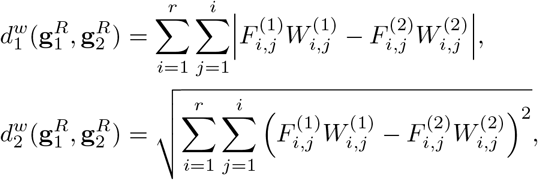

where **W**^(1)^ and **W**^(2)^ are the weight matrices associated with 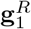 and 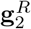, respectively. Figure S3 shows the weight matrix **W** associated with the example heterochronous ranked genealogy and its **F**-matrix in Figures S2(C) and (D).

#### Proposition 2.

The weighted distances 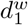 and 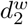 are metrics.

The proof can be found in Section S3. Our distances on ranked tree shapes and ranked genealogies are distances between trees with the same number of leaves and, in the heterochronous case, the same number of leaves and the same number of sampling events. The extension to cases in which the numbers of sampling events differ but the total numbers of leaves remain the same is described in the Materials and Methods section.

In the following section, we propose sample summary statistics based on our metrics *d*_1_ and *d*_2_ for ranked tree shapes and 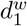 and 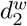 for ranked genealogies.

### Ranked tree shape summary statistics

We use our proposed distances to define a notion of mean value and dispersion value from a finite sample 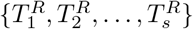 of ranked tree shapes with *n* leaves.

Our proposed measures of centrality are the *L*_2_-medoid sets defined as:

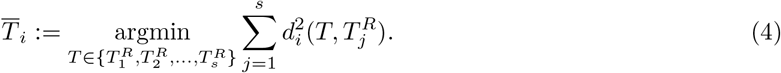

for *i* = 1, 2. We note that our definition of the *L*_2_-medoid set corresponds to the ranked tree shapes in the sample that minimizes the sum of squared distances as opposed to minimize the sum of distances. In addition, when the sample is replaced by the complete population of ranked tree shapes with *n* leaves or when we allow the *L*_2_-medoid to belong to the population of trees but not necessarily to be a sampled tree, Equation 4 corresponds to the *Fréchet mean* or *barycenter* under uniform sampling probabilities (Bacák, 2014).

We use the following as a measure of dispersion around the medoid for *i* = 1, 2:

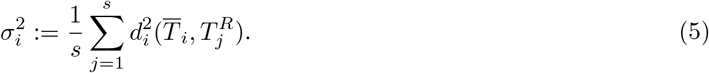

Similarly, given a finite sample of ranked genealogies 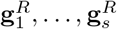 with *n* leaves, the *L*_2_-medoid set is defined as:

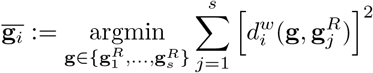

for *i* = 1, 2. Similarly, our empirical measure of dispersion for ranked genealogies is the average sum of squared distances to the medoid.

### Adapting other tree metrics to ranked tree shapes and ranked genealogies

#### Other distances on ranked tree shapes

Although there are no other metrics or distances on ranked tree shapes, we can adapt other tree distances to the space of ranked tree shapes and compare them to our metric. We start with two adaptations of metrics that are originally defined on the space of labeled genealogies: the BHV distance and the KC distance.

#### The Billera-Holmes-Vogtmann metric (BHV) metric

The BHV space (Billera et al., 2001) is obtained by representing each labeled genealogy **g**^*L*^ with edge set *ε* by a vector in the Euclidean orthant 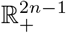, whose coordinates correspond to edge lengths. The BHV space is the union of (2*n −* 3)!! orthants. The BHV distance (*d*_BHV_) between two labeled genealogies is defined as a geodesic, the shortest path connecting two points that lies inside the BHV space. Note that unranked and labeled genealogies with positive intervals between coalescent and sample times are effectively ranked and labeled genealogies. To adapt the BHV distance to the space of ranked tree shapes, we define a modified BHV metric, *d*_BHV-RTS_ as follows:

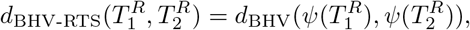

where *ψ* maps a ranked tree shape to its corresponding ranked labeled genealogy by assigning a uniquely defined label to each leaf and assigning a unit length to each change point time interval (*u_i_, u_i−_*_1_).

The unique assignment of the leaf labels *ℓ*_1_, …, *ℓ_n_* on a ranked tree shape consists in assigning labels in increasing index order starting with leaves subtending from nodes closer to the bottom and ending with leaves subtending closer to the root. Detailed description of the unique label assignment can be found in Materials and Methods.

We could alternatively define another BHV-based distance *d*_BHV-RTS*_ on ranked tree shapes as follows:

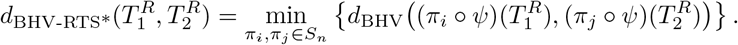

Here, *ψ* assigns an initial labeling to a ranked tree shape and assigns unit branch lengths. *S_n_* is the set of all permutations of the set of leaf labels *{ℓ*_1_, …, *ℓ_n_}*. Although *d*_BHV-RTS*_ is a valid distance on ranked tree shapes and perhaps more natural than *d*_BHV-RTS_, it requires computing BHV distances between all possible pairs of permutations of leaf labels of the two ranked tree shapes. The number of such possible pairs is exponential in the number of leaves and hence it becomes computationally intractable for large *n*. In our results, we only analyze *d*_BHV-RTS*_ for the *n* = 5 case.

#### The Kendall-Colijn (KC) metric

The KC metric is another metric on labeled genealogies (Kendall and Colijn, 2016). A labeled genealogy 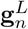 with *n* leaves is represented by an 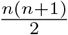-dimensional vector 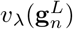 that is a convex combination of two vectors: 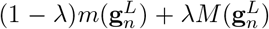, is a vector that concatenates *n* repetitions of one and a vector whose entry corresponds to the number of edges between the most recent common ancestor of a pair of leaves and the root; 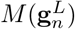 is a vector that concatenates the vector of leaf branch lengths and the branch length between the most recent common ancestor of a pair of tips and the root.

The KC distance *d*_*KCλ*_ with *λ* > 0 between two labeled genealogies is the Euclidean norm of the difference between the two KC vector representations of the labeled genealogies. When *λ* = 0, the KC distance becomes a distance on the space of labeled unranked topologies: *d*_KC,0_.

To adapt the KC distance to the space of ranked tree shapes, we propose two distances. We first define a KC-based distance on ranked tree shapes, *d*_KC-RTS_, as follows

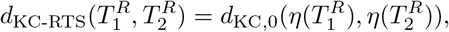

where *η* maps a ranked tree shape to a labeled unranked tree shape by removing internal node rankings and uniquely labeling leaves following the procedure described for *d*_BHV-RTS_.

Alternatively, we can adapt the KC distance on labeled genealogies to define *d*_KC-RTS*_ by uniquely labeling the tips of the ranked tree shape, artificially assigning change point intervals length 1 as in *d*_BHV-RTS_, and *λ* = 0.5 to account for rankings, as follows:

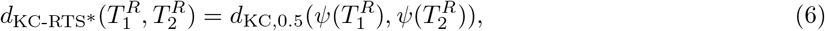

where *ψ* is the mapping previously defined for *d*_BHV-RTS_.

#### The Colijn-Plazzotta (CP) metric

The CP metric is defined on tree shapes (Colijn and Plazzotta, 2018). The CP metric *d*_CP_ is defined as the Euclidean norm (*L*_2_-norm) of the difference between two vectors that uniquely describe the two tree shapes. Each node of a tree is labeled by an integer recursively from tips to the root. The *i*th entry of the CP vector representing a tree shape records the frequency of the tree nodes labeled with integer *i*. We define a modified CP distance *d*_CP-RTS_ on ranked tree shapes as

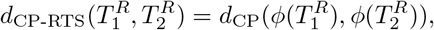

where *ϕ* returns the corresponding tree shape of a ranked tree shape by removing the labels of its internal nodes. We note that *d*_CP-RTS_ is not a metric but a pseudometric: all pairs of different ranked tree shapes with the same shape will have *d*_CP-RTS_ = 0. In addition, *d*_CP-RTS_ does not account for heterochronous sampling, so we exclude *d*_CP-RTS_ from our analyses on heterochronous ranked tree shapes.

#### The Robinson-Foulds (RF) distance

The RF distance on labeled and unranked tree shapes (Robinson and Foulds, 1981) is also a popular metric that can be adapted to ranked tree shapes by labeling the leaves with our unique labeling scheme: 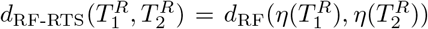, where *η* is the map defined for *d*_KC-RTS_.

#### Other distances on ranked genealogies

We now present the modified BHV and KC distances so that they can be compared to our proposed distances on ranked genealogies.

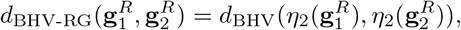

where *η*_2_ maps a ranked genealogy to a labeled ranked genealogy by uniquely labeling leaves as described for *d*_BHV-RTS_. Similarly,

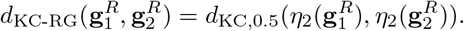

## Results

Having introduced the metrics on the space of ranked tree shapes and the space of ranked genealogies, we examine the behavior of our metrics in a schematic example for *n* = 5, simulated data under various models, and in real human influenza A virus data. The details of simulation and computation steps can be found in the Materials and Methods section.

### Interpreting proposed distances between ranked tree shapes of *n* = 5 leaves

We first compare our distances on ranked tree shapes of *n* = 5 leaves with our adaptations of other metrics. In Figure 4, we show all 5 possible ranked tree shapes and their corresponding pairwise *d*_1_ distances. Trees 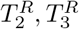 and 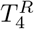 are trees with the same tree shape but different ranked tree shapes. The pairs 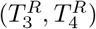 and 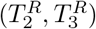 differ by one ranking switch between two cherries—ranking 2 by 3 and ranking 3 by 4, respectively— and their pairwise distance is *d*_1_ = 1. The *d*_1_ distance between the pair 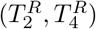 is 2. Indeed, to go from 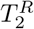 to 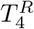, we need two ranking switches: ranking 2 by 3 and then ranking 3 by 4. All pairwise distances are shown in Figure 5. Qualitative behaviors of distances *d*_KC-RTS_ and *d*_BHV-RTS*_ between the trees 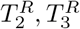 and 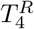 mirror *d*_1_.

**Figure 5:**
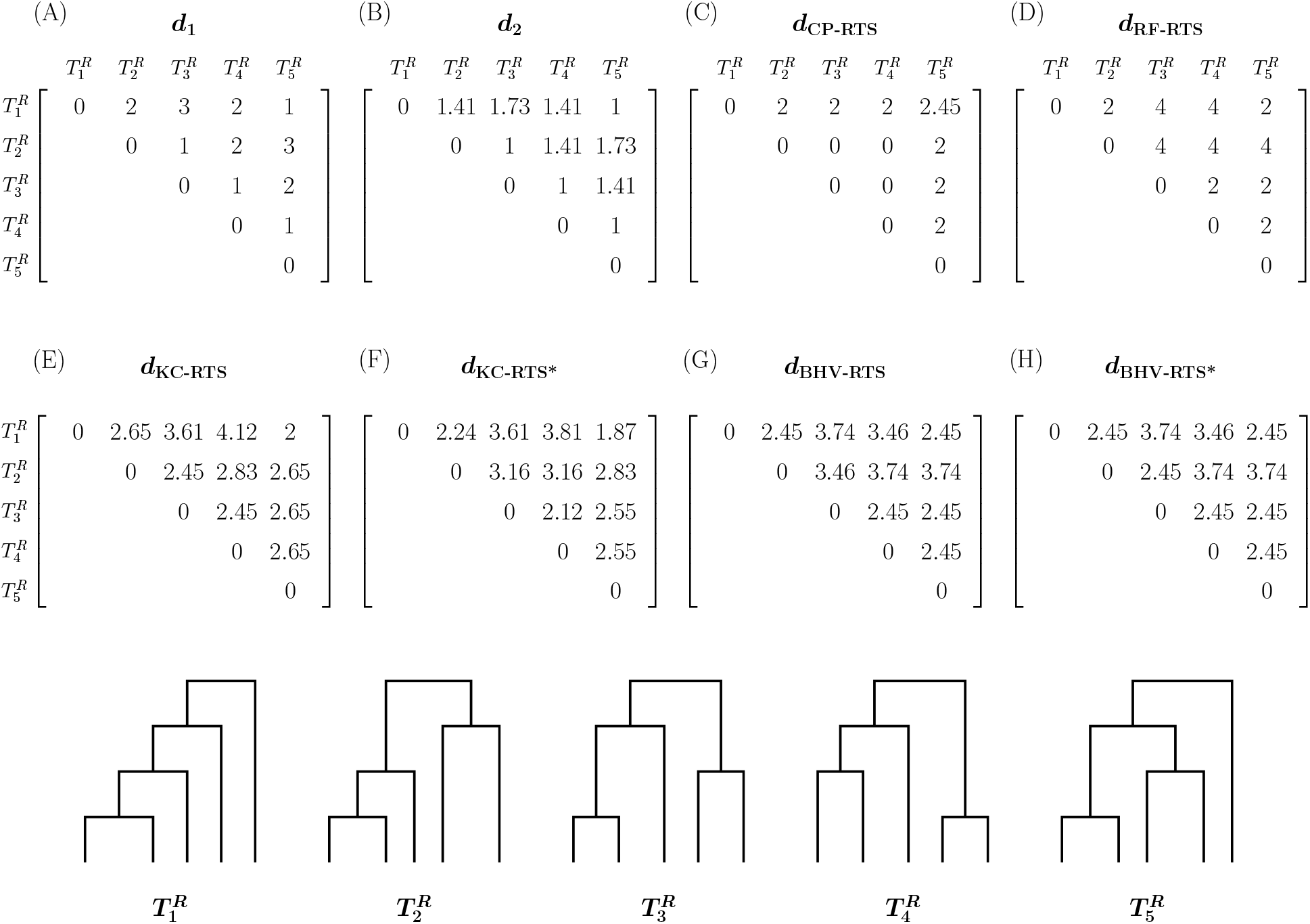
Comparisons of metrics applied to isochronous ranked tree shapes with *n* = 5.

The pairs 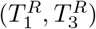 and 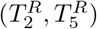 have the maximum *d*_1_ distance. In both cases, we need a ranking switch and a change of tree shapes to move from one tree to the other in each pair. The pairs of trees with the maximum *d*_BHV-RTS_ or *d*_BHV-RTS*_ distance are the same two pairs with the maximum *d*_1_ distance. However, an additional pair, 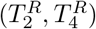, also has the same *d*_BHV-RTS_ and *d*_BHV-RTS*_ maximum value even when 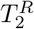 and 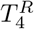 differ by two ranking change events but have the same tree shape. Unlike *d*_BHV-RTS_ or *d*_BHV-RTS*_, our *d*_1_ distance penalizes ranking changes and tree shape changes differently. The distances between pairs 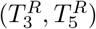 and 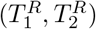 are penalized more with the *d*_1_ distance than with the *d*_BHV-RTS*_ since it involves changes in tree shapes.

In the analysis of the *n* = 5 case, our *d*_1_ distance is more similar to *d*_BHV-RTS*_ than to any other of the distances considered. The second most similar distance to *d*_1_ is *d*_RF-RTS_. In general, we notice that *d*_BHV-RTS*_ penalizes changes in internal node rankings and changes in tree shapes equally, whereas our *d*_1_ distance penalizes tree shapes changes more than ranking changes. As mentioned previously, computation of *d*_BHV-RTS*_ is expensive; however, *d*_BHV-RTS_ and *d*_BHV-RTS*_ produce the same pairwise distances except for the 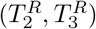 pair.

In the following results sections, we use multidimensional scaling (MDS) (Mardia, 1978) to embed our distance metrics into Euclidean spaces of two dimensions for ease of visualization and interpretation. We set the two axes of each two-dimensional MDS plot to have the same scale for correct interpretations (Holmes and Huber, 2019). The MDS representation of trees is useful for exploring large sets of trees, but it has also been recognized to have some limitations (Hillis et al., 2005; Chakerian and Holmes, 2012). For this reason, in addition to MDS plots, we include additional summaries to assess the behavior of our metrics. Initially, we assume that our trees are isochronous, that is, all samples are obtained at time 0.

### Separation between ranked tree shapes with different distributions

To assess how our proposed metrics differentiate ranked tree shapes sampled from distributions with different degrees of balance, we simulated 1000 ranked tree shapes with *n* = 100 leaves under the beta-splitting model for each *β*-value in *{−*1.9, *−*1.5, *−*1, 0, 100} representing a sequence from unbalanced to balanced. From the resulting 5000 total simulated ranked tree shapes, we computed the 5000 *×* 5000 pairwise distance matrices with *d*_1_, *d*_2_, *d*_BHV-RTS_, *d*_CP-RTS_ and *d*_KC-RTS_ distances. The MDS plots are depicted in Figure 6. Each dot corresponds to a tree, and each color corresponds to the sampling distribution for a specified *β* value. Figure 6(C) shows the *L*_2_-medoid trees using our *d*_1_ distance. The total distance explained by the first two MDS dimensions are 91.4% using *d*_1_ (Figure 6(A)) and 94.4% using *d*_2_ (Figure 6(B)). The 2-dimensional MDS mappings using the other distances explain less than 80%. Our distances visually discriminate trees with different balance distributions to a greater extent than *d*_BHV-RTS_ and *d*_KC-RTS_; *d*_CP-RTS_ shows similar performance.

**Figure 6:**
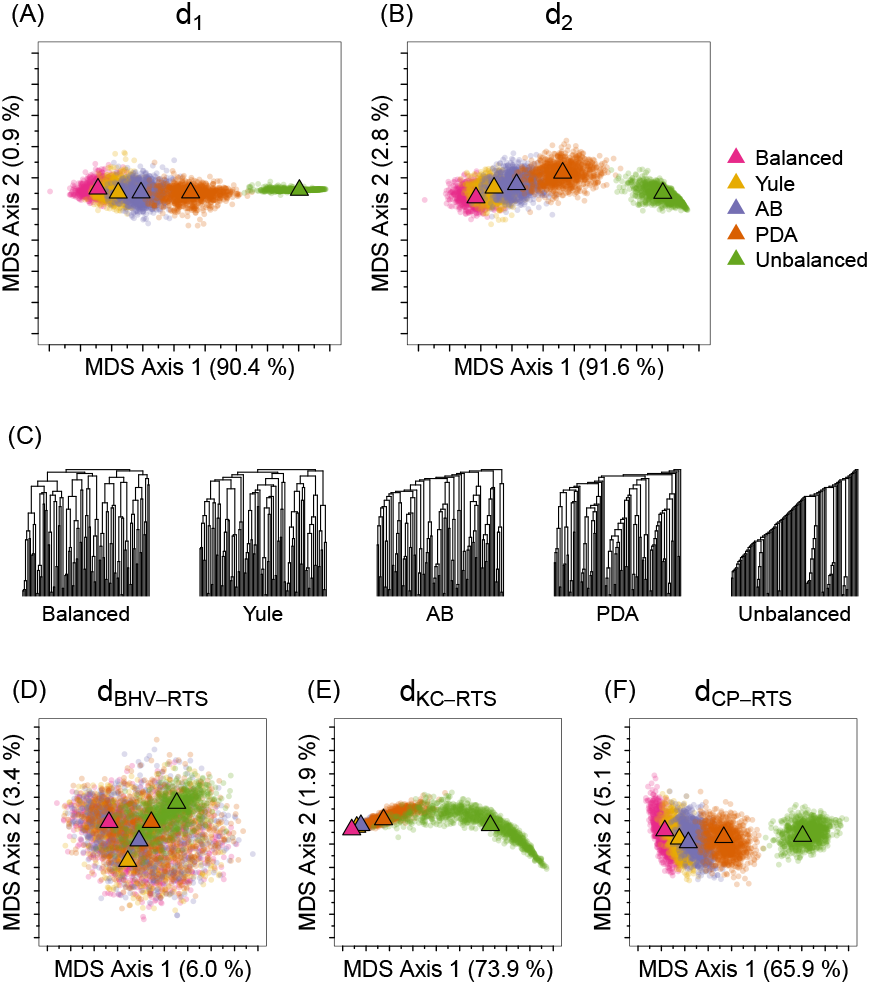
MDS representation of distances between 5000 simulated isochronous ranked tree shapes of *n* = 100 leaves, aggregated from five different beta-splitting models. 1000 isochronous ranked tree shapes were simulated from each of the following models: the balanced model (*β* = 100), the Yule model (*β* = 0), the Aldous branching (AB) model (*β* = *−*1), the proportional-to-distinguishable-arrangements (PDA) model (*β* = *−*1.5), and the unbalanced model (*β* = *−*1.9). (A) MDS of the *d*_1_ metric. (B) MDS of the *d*_2_ metric. (C) *L*_2_-medoid trees from each distribution using the *d*_1_ metric. MDS plots for (D) *d*_BHV-RTS_, (E) *d*_KC-RTS_, and (F) *d*_CP-RTS_. In each MDS plot, the triangle represents the *L*_2_-medoid tree of 1000 points for a specified model.

In Figure S5, we show the confusion matrices for the trees plotted in Figure 6. For *i, j ∈ {*1, *…,* 5}, each (*i, j*)-th entry of the matrix represents the percentage of trees simulated from the *i*-th distribution that are closest to the *L*_2_-medoid of the *j*-th distribution, where closeness is measured by the distance function. As an indication of overall separation, we average the diagonal entries of the confusion matrices. A larger value indicates better separation of the corresponding distance. Matrices in Figure S5 confirm the greater discrimination for *d*_1_ and *d*_2_: about 83% of the trees are closest to the *L*_2_-medoid of their originating distribution with *d*_1_ and *d*_2_ distances, 70.5% with *d*_KC-RTS_, 75.0% with *d*_CP-RTS_, and only 20.3% with *d*BHV-RTS.

Table S1(A) shows the dispersion statistic (Section S4.1) for each tree distribution and for each distance function. The group of unbalanced trees (green in Figure 6) is the group with the least dispersion according to *d*_1_, *d*_2_, and *d*_BHV-RTS_ but not according to the other three metrics. The MDS figures (Figure 6) accord with Table S1: both distances *d*_KC-RTS_ and *d*_CP-RTS_ show a linear trend that increases dispersion with degrees of unbalancedness. This pattern is particularly evident in Figure 6(E).

We next simulated trees from the alpha-beta splitting model (Maliet et al., 2018). We generated 1000 random trees with *n* = 100 for each *α* in *{−*2, *−*1, 0, 1, 2}, producing 5000 total trees. By varying the value of *α* while keeping the balance parameter (*β*) fixed, we test the performance of our metrics in distinguishing between ranked tree shapes with small family sizes closer to the root (*α* < 0) and ranked tree shapes with small family sizes closer to the tips (*α* > 0). Our two-dimensional MDS representations are shown in Figure 7 and their corresponding confusion matrices in Figure S6. Our metrics, *d*_1_ (Figure 7(A)) and *d*_2_ (Figure 7(B)), when compared to the other three metrics, distinguish to a greater extent between trees with positive and negative *α* parameters. More than 75% of the trees are closest to the *L*_2_-medoid of their originating distributions with *d*_1_ and *d*_2_, and less than 35% have this property with the other distances.

**Figure 7:**
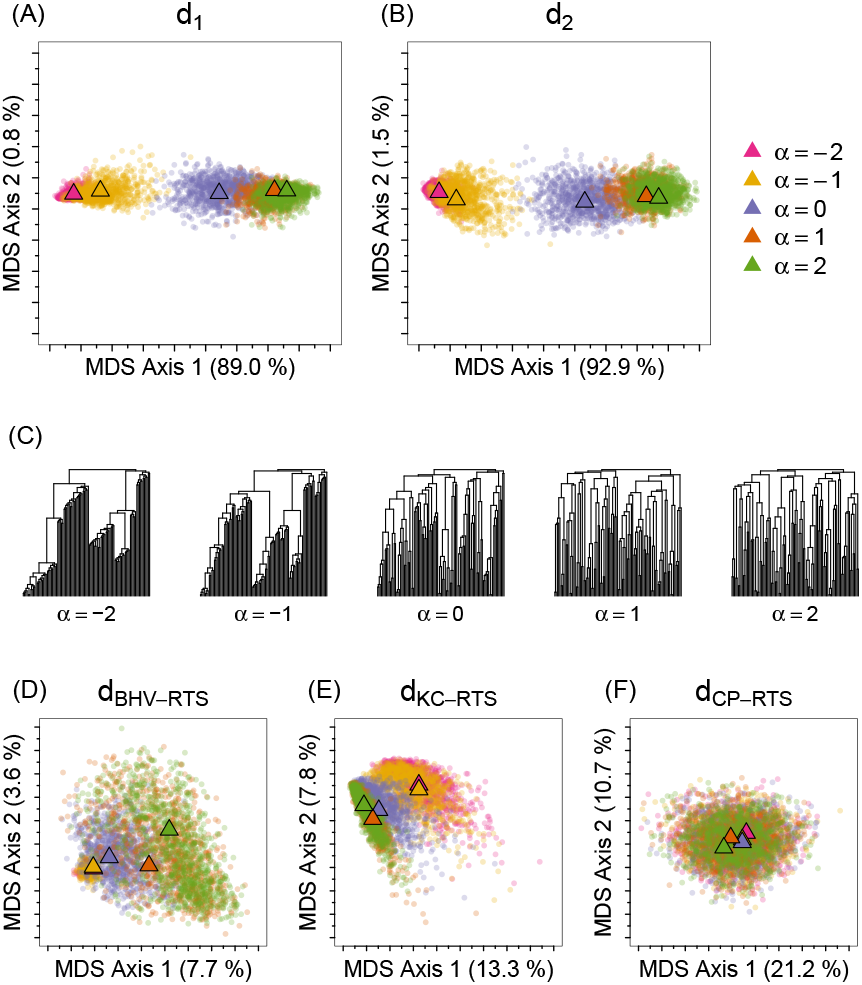
MDS representation of distances between 5000 simulated isochronous ranked tree shapes of *n* = 100 leaves, aggregated from five different alpha-beta splitting models. 1000 isochronous ranked tree shapes were simulated for each of *α* values in 2, 1, 0, 1, 2 . Different *α* generates different distributions of internal node ranking while keeping the same tree shape distribution at *β* = 0. (A) MDS of the *d*_1_ metric. (B) MDS of the *d*_2_ metric. (C) *L*_2_-medoid trees from each distribution using the *d*_1_ metric. MDS plots for (D) *d*_BHV-RTS_, (E) *d*_KC-RTS_, and (F) *d*_CP-RTS_. In each MDS plot, the triangle represents the *L*_2_-medoid tree of 1000 points for a specified model.

The first two dimensions in MDS explain more than 90% of the total *d*_1_ and *d*_2_ distances, whereas the other metrics explain less than 35%. Although the MDS visualizations in Figures 7(A) and (B) show that most clusters of trees are well separated according to their sampling distributions with *d*_1_ and *d*_2_, a large overlap exists between groups of trees with *α* = 1 and *α* = 2. The observed similarity is evident from the *L*_2_-medoid trees (Figure 7C). Figure S6 confirms these conclusions obtained by visually inspecting the MDS plots of Figure 7. We note that *d*_KC-RTS_ has better performance than *d*_BHV-RTS_ and *d*_CP-RTS_ in discriminating simulated trees from the alpha-beta distributions with different *α* values.

The dispersion statistics in Table S1(B) indicate that trees from the alpha-beta splitting model with *α* = 0 are the most dispersed group according to our *d*_1_ and *d*_2_ metrics. This, however, is not the case with the other distances. In particular, no variations in dispersion across different distributions are observed using *d*_CP-RTS_. *d*_BHV-RTS_ shows a correlated trend between *α* and the dispersion of the corresponding distribution: trees from a distribution with large *α* have higher dispersion than trees with small *α*.

Our results confirm that our metrics capture both shape and rankings more effectively than the other adapted metrics on ranked tree shapes in that simulation models differing in these features are more easily discriminated with our distances than with the others. In addition, our *d*_1_ and *d*_2_ distances show good embedding in two-dimensional MDS, with a high proportion of the total distance explained.

### Separation between genealogies with different branch length distributions

When a population undergoes changes in population size *λ*(*t*), the branch length distribution of the ranked genealogies depends on the population history *λ*(*t*) as shown in Eq. 3. Under the neutral coalescent, the tree topology distribution in Eq. 1 is independent of the branch lengths. To investigate the performance of our weighted metrics 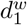 or 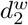 on genealogies in separating trees according to their branch length distribution, we generated 1000 random ranked genealogies with *n* = 100 leaves from the Tajima coalescent with each of the following demographic scenarios:

1. Constant effective population size: *λ*(*t*) = *N*_0_;
2. Exponential growth: *λ*(*t*) = *N*_0_*e*^*−*0.01*t*^;
3. Seasonal logistic trajectory:

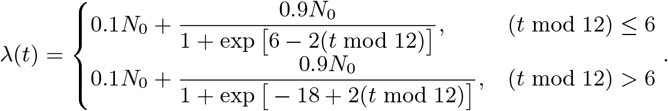

The coalescent trees under each specified population trajectory were simulated with *N*_0_ = 10000. The functional form chosen and parameter values for the seasonal logistic trajectory mimic estimated trajectories of Human Influenza A virus in temperate regions (Vijaykrishna et al., 2015). We removed the leaf labels and retained branch lengths to produce isochronous ranked genealogies. We produced 3000 *×* 3000 pairwise distance matrices using the weighted metrics 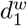 and 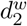.

Figures 8(A) and (B) show that our weighted metrics distinguish the three different demographic scenarios. The two-dimensional MDS accounts for 72.1% of the total distance, with the first MDS coordinate separating the uniform trajectory from the others, and the second coordinate distinguishing the exponential trajectory from the logistic trajectory.

**Figure 8:**
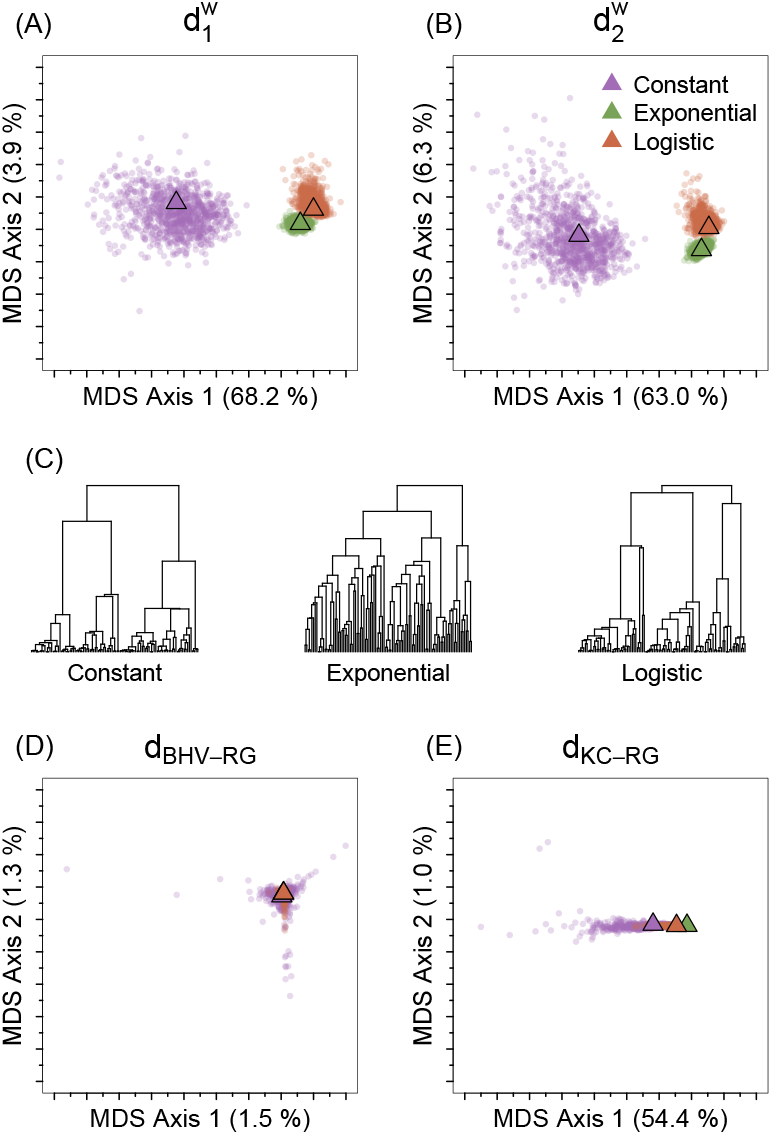
MDS representation of distances between 3000 simulated isochronous ranked genealogies of *n* = 100 leaves under different demographic models. The number of simulated genealogies per population model is 1000. (A) 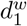 metric, (B) 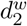 metric, (C) *L*_2_-medoid genealogies from each distribution using the 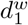 metric, (D) *d*_BHV-RG_, and (E) *d*KC-RG.

A comparison of panels within Figure 8 shows that *d*_BHV-RG_ is far less informative than any of our metrics. A cline differentiating the three distributions is somewhat present using *d*_KC-RG_ (Figure 8(E)), far less so than in Figures 8(A) and (B). Our metrics’ ability to distinguish different ranked genealogy distributions is also apparent in the confusion matrix in Figure S7, where our metrics display near perfect performance. Across all three distributions, 99.8% of the trees are closest to the *L*_2_-medoid of their originating distribution with 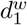 and 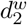; corresponding values are 33.4% with *d*_BHV-RG_ and 74.0% with *d*_KC-RG_.

### Separation between heterochronous ranked tree shapes with different sampling events

To demonstrate that our metrics are sensitive to sampling schedules, we simulated trees with *n* = 100 leaves under heterochronous sampling with two sampling scenarios. For the first set of trees, we selected 90 samples at time 0 and the remaining 10 samples at distinct times with sampling times drawn uniformly at random from (0, 10^4^], i.e., **n**= (90, 1, 1, 1, 1, 1, 1, 1, 1, 1, 1). In the second set of trees, 80 samples were drawn at time 0 and the remaining 20 were sampled in pairs at ten distinct random sampling times drawn uniformly from (0, 10^4^], i.e., **n**= (80, 2, 2, 2, 2, 2, 2, 2, 2, 2, 2). We generated 1000 coalescent trees per sampling scheme assuming a constant effective population size trajectory of *N*_0_ = 10000. We then removed leaf labels to produce the 2000 *×* 2000 distance matrices with all applicable distances.

The resulting MDS visualizations are displayed in Figures 9. Our metrics and *d*_BHV-RTS_ show a clear separation between the two distributions along the first two MDS axes. The total distance explained in the two-dimensional space is higher using our metrics, 66.5% and 76.4% for *d*_1_ and *d*_2_, respectively, compared to 13.5% of *d*_BHV-RTS_. *d*_KC-RTS_ and *d*_CP-RTS_ exhibit less discrimination than the other three distances. The confusion matrices in Figure S8 confirm that our metrics distinguish different sampling schemes better than the other three metrics compared.

**Figure 9:**
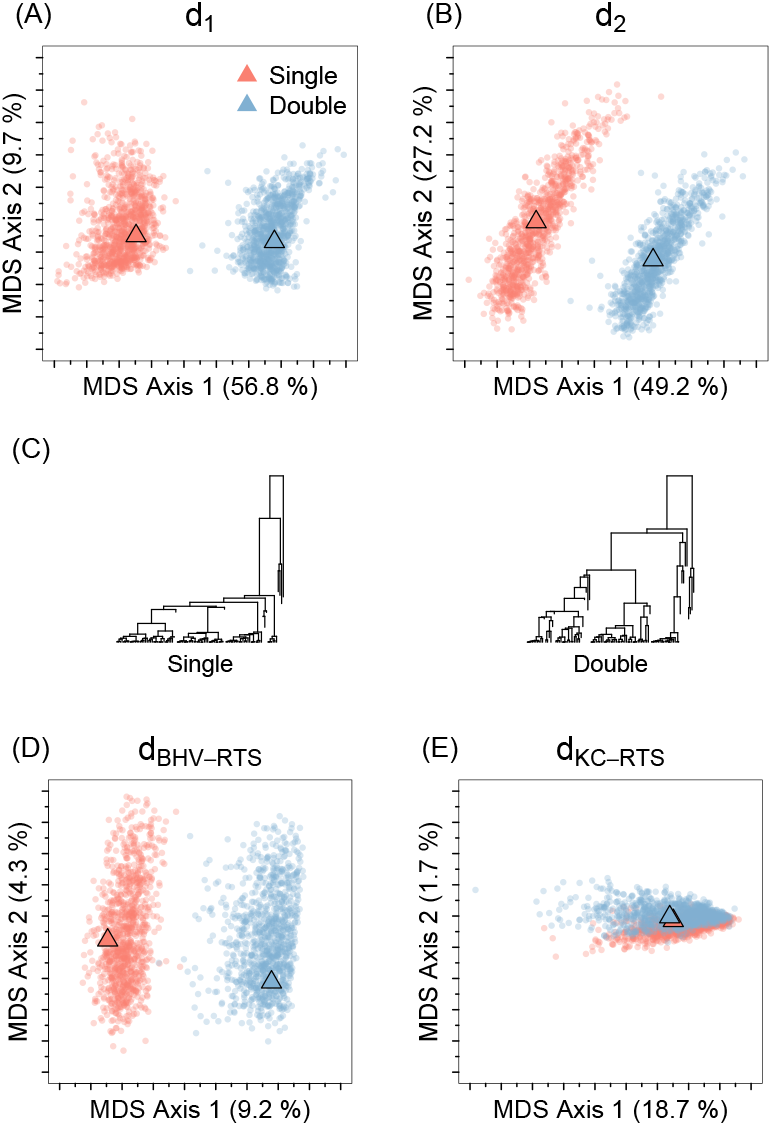
MDS representation of distances between 2000 simulated heterochronous ranked tree shapes of *n* = 100 with different sampling events. In the “Single” distribution, a single sample is drawn at each of the ten sampling events after the initial sampling event at time 0 with 90 samples: **n**= (90, 1, 1, 1, 1, 1, 1, 1, 1, 1, 1). In the “Double” distribution, a pair of samples is drawn at each of the ten sampling events after the initial sampling event at time 0 with 80 samples: **n** = (80, 2, 2, 2, 2, 2, 2, 2, 2, 2, 2). The sampling times of the ten sampling events after *t* = 0 are selected uniformly at random from (0, 10^4^] in both distributions. The resulting simulated trees with *n* = 100 taxa are from the neutral coalescent model with constant effective population size of 104. (A) *d*_1_ metric, (B) *d*_2_ metric, (C) *L*_2_-medoid trees from each distribution using the *d*_1_ metric, (D) *d*_BHV-RTS_, (E) *d*_KC-RTS_, and (F) *d*CP-RTS.

### Application to simulated influenza data

Before applying our metric to human influenza data, we examined our metric on simulated heterochronous genealogies that mimic hypothesized human influenza population size trajectories in temperate and tropical regions. In temperate climates, such as New York, influenza epidemics display seasonality, with peaks occurring during the winter months, whereas in regions with tropical or subtropical climates, influenza activity is persistent throughout the year (Tamerius et al., 2013).

For our hypothesized temperate region influenza dynamics, we assumed the seasonal logistic trajectory with *N*_0_ = 100. We adopted preferential sampling (Karcher et al., 2016) to model probabilistic dependence of sampling times on effective population size by drawing sampling times from an inhomogeneous Poisson process with intensity proportional to the effective population size. The lower and upper sampling time bounds were set to 0 and 48, respectively. The resulting sampling event had *n* = 104 total samples. Parameter values were chosen following (Karcher et al., 2016) to mimic realistic Influenza A trajectories. Given the generated sampling event vectors **s** and **n**, we generated 1000 random coalescent trees for the temperate region.

In tropical regions, influenza activity is more stable throughout the year and hence, we assumed a constant trajectory with *N*_0_ = 100. We simulated 1000 random coalescent trees with sampling times of *n* = 104 samples randomly drawn from a uniform-[0, 48] distribution.

We constructed 2000 *×* 2000 distance matrices with each of the metrics applicable for ranked genealogies: 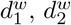, *d*_BHV-RG_, and *d*_KC-RG_. Figure 10 shows that the first two MDS components with our metrics 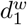 and 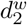 explain 34.7% and 39.2% of the total distance, respectively, with the first component clearly separating between seasonal and tropical distributions. *d*_BHV-RG_ do not distinguish the models to the same extent.

**Figure 10:**
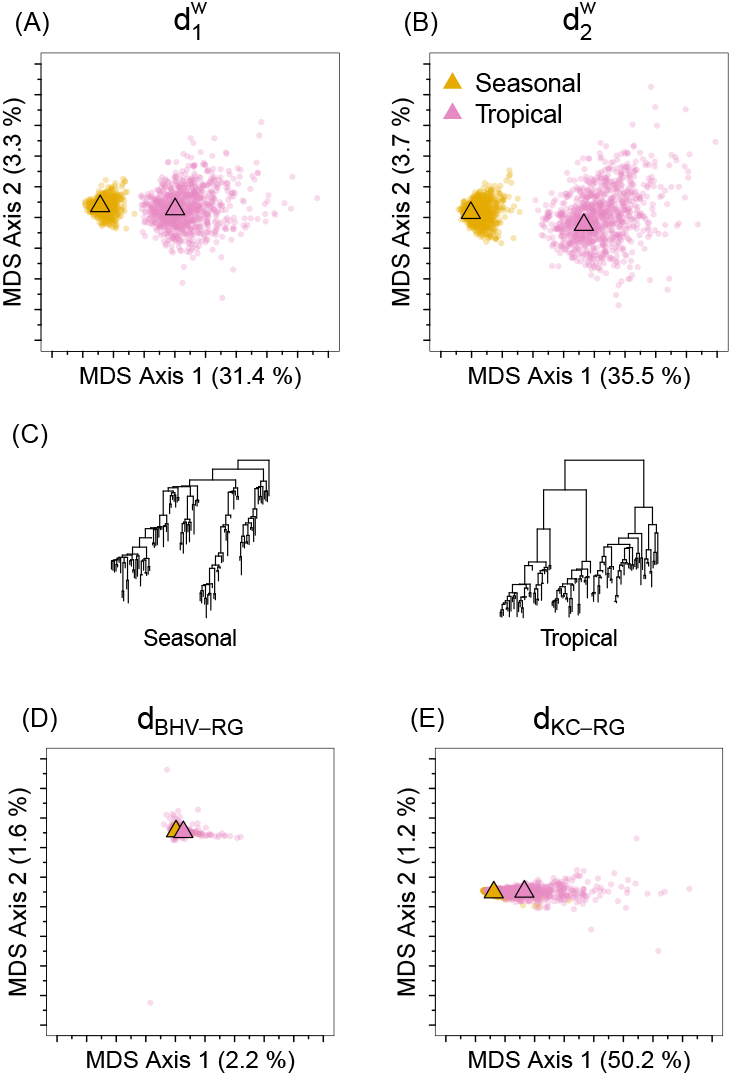
MDS representation of distances between 2000 simulated heterochronous ranked genealogies of *n* = 104 leaves under different demographic models and sampling events. The “Seasonal” distribution represents the population dynamics of influenza in the temperate region, which is modeled with logistic trajectory and preferential sampling reflecting the seasonality of the influenza dynamics. The “Tropical” distribution represents the population dynamics of influenza in the tropical region, which is modeled with a constant population trajectory and uniform sampling. (A) 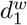 metric, (B) 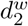 metric, (C) *L*_2_-medoid trees from each distribution using the 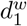 metric, (D) *d*BHV-RG, and (E) *d*KC-RG.

Although the *d*_KC-RG_ metric shows greater variation in the first MDS axis than with our metrics, *d*_KC-RG_ does not distinguish between the two distributions. The lack of separation explained by the first MDS axis in Figure 10(E) can be attributed to the large dispersion in the sampling-time distribution of the tropical-region simulated genealogies. This claim is supported by Figure S9, where ranked genealogies from the tropical distribution are approximately equally likely to be close to the *L*_2_-medoid points of the seasonal and tropical distributions with *d*_KC-RG_. By contrast, our metrics show near perfect diagonal entries. In particular, under the 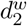 metric, every tree is closer to the *L*_2_-medoid tree of its own distribution than to the other *L*_2_-medoid trees. Across two distributions, 99.7% of the trees are closest to the *L*_2_-medoid of their originating distribution with 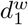 and 100% with 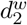. Corresponding numbers are 51.0% with *d*_BHV-RG_ and 74.1% with *d*_KC-RG_.

Table S2(B) shows the dispersion statistics: as expected, all distances reflect higher dispersion in the tropical data than the dispersion observed in the seasonal data.

### Analysis of human influenza A virus from different continents

We next apply our metrics to ranked tree shapes and ranked genealogies sampled from the posterior distributions of human influenza A/H3N2 genealogies from 3 geographic regions—New York, Chile, and Singapore— collected from January 2014 to January 2019 in the Global Initiative on Sharing All Influenza Data’s (GISAID) EpiFlu database. The frequency of the collected samples by collection date and the estimated effective population size trajectories with the BEAST Bayesian Skygrid method are displayed in Figure S11 in Supplementary Material.

For each of the three regions, we sampled 1000 trees from the posterior distribution with BEAST. We then computed our unweighted and weighted distance matrices of 3000 *×* 3000 pairwise distances. Figures 11(A) and (B) show the corresponding MDS plots of the ranked tree shapes using our *d*_1_ and *d*_2_ metrics. All three distributions are well separated in the two-dimensional MDS plots. The first MDS axis separates Singapore from the rest, and the second MDS axis splits Chile from the other two ranked tree shape distributions. Figures 11(A) and (B) show that the three tree distributions have varying degrees of dispersion. The trees from Singapore are more clustered together than the trees from the other two regions. Similarly, the trees from New York are more clustered together than the trees from Chile. The dispersion statistics in Table S2(C) quantify the observed differences in variation among different tree posterior distributions.

**Figure 11:**
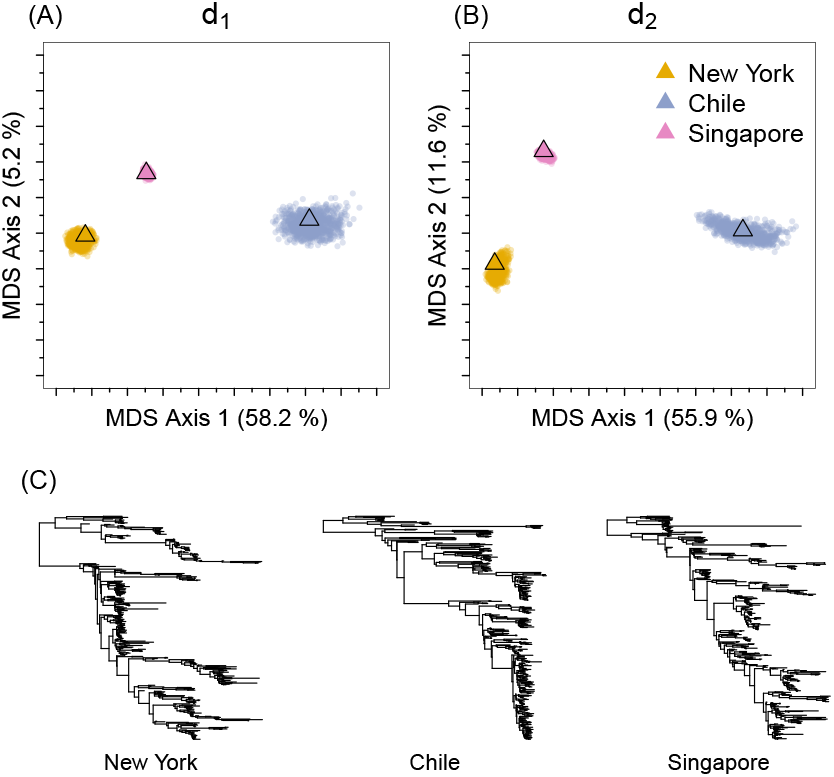
MDS representation of distances between 3000 heterochronous ranked tree shapes sampled from three posterior distributions (1000 trees from each distribution). Observed data consist of 410 sequences of human influenza A virus from each geographic region: Singapore, New York and Chile. (A) *d*_1_ metric, (B) *d*_2_ metric, and (C) *L*_2_-medoid trees from each distribution using the *d*_1_ metric.

The same patterns of dispersion are observed with the weighted metrics 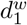 and 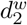 of ranked genealogies (Figure 12). The dispersion observed in the MDS plots reflects the high variance in branch lengths.

**Figure 12:**
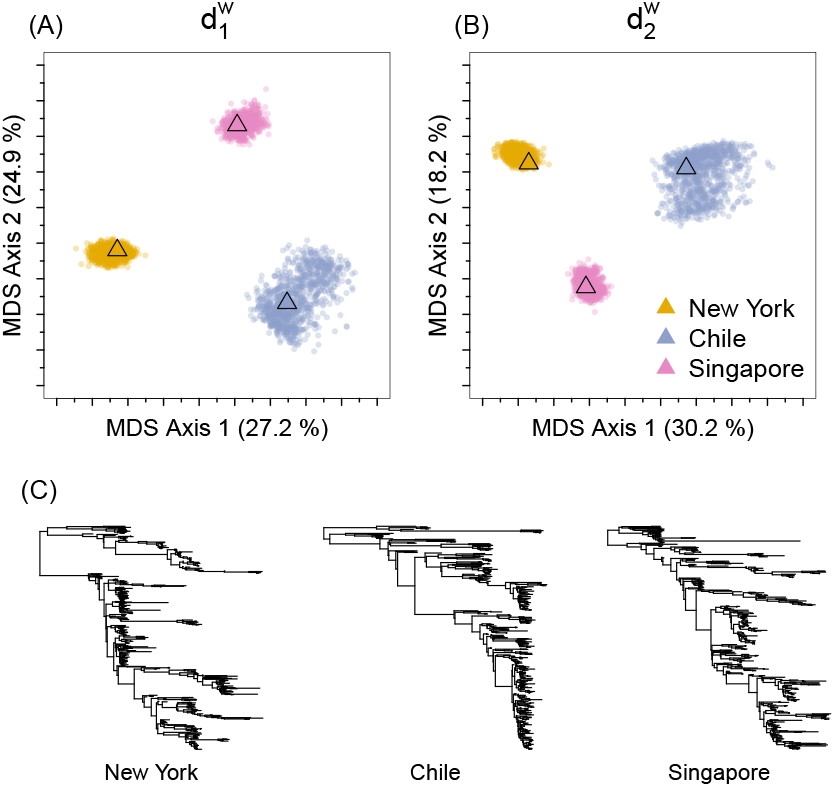
MDS representation of distances between 3000 heterochronous ranked genealogies sampled from three posterior distributions (1000 trees from each distribu-tion). Observed data consist of 410 sequences of human influenza A virus from each geographic region: Singapore, New York and Chile. (A) 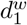 metric, (B) 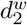 metric, and (C) *L*_2_-medoid trees from each distribution using the 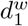 metric.

## Discussion

Ranked tree shapes and ranked genealogies are used to model the dependencies between samples of molecular data in the fields of population genetics and phylogenetics. These tree structures have a particular value in studying heterochronous data and in providing a resolution that is fine enough to be informative but is computationally more feasible than finer resolutions. Although many other distances are defined on trees, they are primarily defined on spaces of labeled tree topologies or tree-shape structures without ranking of internal nodes. The applications of such distances are limited when comparing detailed hierarchical tree structures from different sets of individuals with little or no overlap.

In this manuscript, we equipped the spaces of ranked tree shapes and ranked genealogies with a metric. By defining a bijection between ranked tree shapes and a triangular matrix of integers, we defined distance functions between ranked tree shapes as the *L*_1_ and *L*_2_ norms of the difference between two matrices. We extended our metrics to heterochronously sampled ranked tree shapes and genealogies. Our distances on ranked genealogies are the *L*_1_ and *L*_2_ norms of the difference between two matrices whose elements are the products of the corresponding entries of the ranked tree shape matrix representation and weights given by the genealogy branch lengths. To apply our distances to real data for comparing ranked genealogies or ranked tree shapes with different sampling events, we proposed an augmentation scheme, increasing the dimension of the matrix representations, and we defined our distances as *L*_1_ and *L*_2_ norms of the difference between extended matrices of the same size.

We compared our distance to projections of the BHV (Billera et al., 2001), Colijn-Plazotta (Colijn and Plazzotta, 2018), and Kendall-Colijn (Kendall and Colijn, 2016) distances on the spaces of ranked tree shapes and genealogies (when applicable). The comparison showed that our distances perform better than the other metrics in separating ranked tree shapes from distributions with different balance parameters (Figure 6), different family size compositions with respect to time (Figure 7), and different branch length distributions (Figure 8).

Our proposed distances do not equally penalize all types of tree differences. The analyses showed that our distances penalize coalescence-type changes—whether the coalescence is between two unlabeled leaves, a leaf and an internal (ranked) branch, or between two internal (ranked) branches— more than ranking changes. Events that change the tree shape incur a higher penalty. In addition, our metric penalizes changes near the root more than near the bottom of the trees. These attributes accord with desirable properties for making biologically meaningful metrics.

We used our distances to summarize the distribution of trees in 2-dimensional MDS representations and propose using the *L*_2_-medoid tree, a sample version of the Fréchet mean, as the central point of a sample of trees. We proposed a sample version of the Fréchet variance to quantify uncertainty. In our simulations, the *L*_2_-medoid trees summarized the center of the distributions, and in some cases, they looked similar to the maximal clade credibility trees. Despite this, however, the geometry of the spaces induced with our metrics is still unknown, and the mathematical properties of such summary statistics require further exploration.

Our metrics provide the basis for a decision-theoretic statistical inference that can be constructed by finding the best-ranked genealogy that minimizes the expected error or loss function, which, in turn, is a function of the tree distance. Further, our proposed metric provides a tool for evaluating convergence and the mixing of Markov chain Monte Carlo procedures on ranked genealogies.

Our proposed distances inherit the properties of *L*_1_ and *L*_2_ distances of symmetric positive definite (SPD) matrices. In fact, our matrix representation can be modified to a SPD matrix by replacing the upper matrix triangle with the transposed entries of the original lower triangular matrix. In this case, our new distance will be twice the original distance and will not change the observed properties. This view of SPD matrix representation of ranked tree shapes makes it possible to explore new distances (Vemulapalli and Jacobs, 2015) on ranked tree shapes and ranked genealogies.

## Materials and Methods

### Metric spaces on heterochronous trees with different numbers of sampling events

We extend our distances to include cases in which the numbers of sampling events differ but the total number of samples remain the same.

Consider two heterochronous ranked tree shapes of *n* leaves, 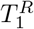 and 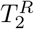, with different numbers of sampling events *m*_1_ and *m*_2_, respectively. In order to compute the distance between 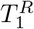 and 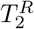 with our metrics, we require the two ranked tree shapes to be represented as **F**-matrices of the same dimension. We accomplish this by inserting artificial sampling events. The detailed steps are as follows. Note that the following formulation is done going backwards in time with time increasing from the present to the past.

For *i* = 1, 2, let 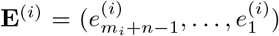 be the vector of ordered sampling and coalescence events where 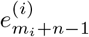 denotes the most recent sample event 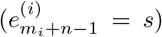 assumed to occur at time 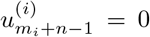. 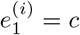 denotes the coalescent event at time 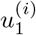 corresponding to the most recent common ancestor of the samples in 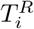. Each 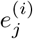 is either a sampling event 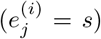 or a coalescent event 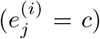. The event of type *c* occurs *n* − 1 times and the event of type *s* occurs *m_i_* times in **E**^(*i*)^. In the example illustrated in Figure S4, the event vectors for 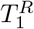 (Figure S4(A)) and 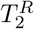 (Figure S4(B)) are **E**^(1)^ = (*s, c, s, c, s, c*) and **E**^(2)^ = (*s, s, s, c, s, c, c*), respectively.

We first align all *n* − 1 coalescent events between the two trees by adding empty spaces when needed as depicted in Figure S4(C). Once all the type-*c* events are aligned, we next align the sampling events between two successive coalescent events or between *t* = 0 and the first *c* event. When aligning the events of type *s* between the two trees, we pair the events of type *s*, starting from the most recent events. The event vector alignment is demonstrated in Figure S4(C). If one tree has more type-*s* events than the other in a given intercoalescent interval, we insert the excess artificial sampling events, denoted by *a*’s, in the tree with fewer type-*s* events in that interval. We assign 0 new samples to type-*a* events. For example, in the tree of Figure S4(A), there is only one sampling event before the first coalescent event, whereas there are three sampling events in the tree of Figure S4(B). In Figure S4(D), we illustrate how the two artificial sampling events are added to the first tree in the first interval. The resulting augmented trees are shown in Figures S4(E) and (F) along with their corresponding **F**-matrix representations in Figures S4(G) and (H). We note that by construction, **F**^(1)^ ≠ **F**^(2)^ in these cases. Hence 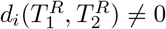 for *i* = 1, 2.

Consider now two heterochronous ranked genealogies of *n* leaves, 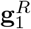 and 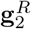, with different number of sampling events *m*_1_ and *m*_2_ respectively. In order to compute the distance between 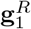 and 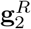 we first augment their **F**-matrix representations as with ranked tree shapes. In addition, we augment their weight matrices *W* ^(1)^ and *W* ^(2)^ by assigning a time to each augmented artificial sampling event. If *n_a_* artificial events are inserted between events 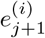 and 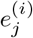, we subdivide the corresponding time interval 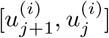 into *n_a_* + 1 intervals with equal length: the times assigned to the *n_a_* augmented artificial events are 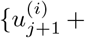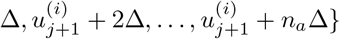, where 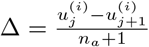.

### Unique labeling scheme of ranked tree shapes

In order to adapt other distances defined on labeled trees to ranked tree shapes, we use the following labeling scheme. We start by labeling the leaves that descend directly from the internal node with the largest rank If the node has two direct descendent leaves with different edge lengths, we label the longer leaf *ℓ*_1_ and the shorter leaf *ℓ*_2_. If the two leaves have the same edge lengths, we label them *ℓ*_1_ and *ℓ*_2_ from left to right. If the node has only one direct descendent leaf, we label it *ℓ*_1_. If there is no descending leaf, no labeling is done. We then move to the node with rank *n −* 1 and continue labeling the leaves by traversing through the internal nodes in descending order of rank until all leaves are labeled. If the current node with rank *k* has two direct descendent leaves and if the last assigned leaf label is *ℓ_j_*, we label the leaf with the longer edge *ℓ_j_*_+1_ and the leaf with shorter edge *ℓ_j_*_+2_; if the leaves have the same edge lengths, we label the pair of leaves *ℓ_j_*_+1_ (left) and *ℓ_j_*_+2_ (right). If the node *k* has only one direct descendent leaf, we label it *ℓ_j_*_+1_. If the node *k* has no direct descendent leaf, no label is assigned. Examples demonstrating our unique labeling scheme are in Figure 13.

**Figure 13:**
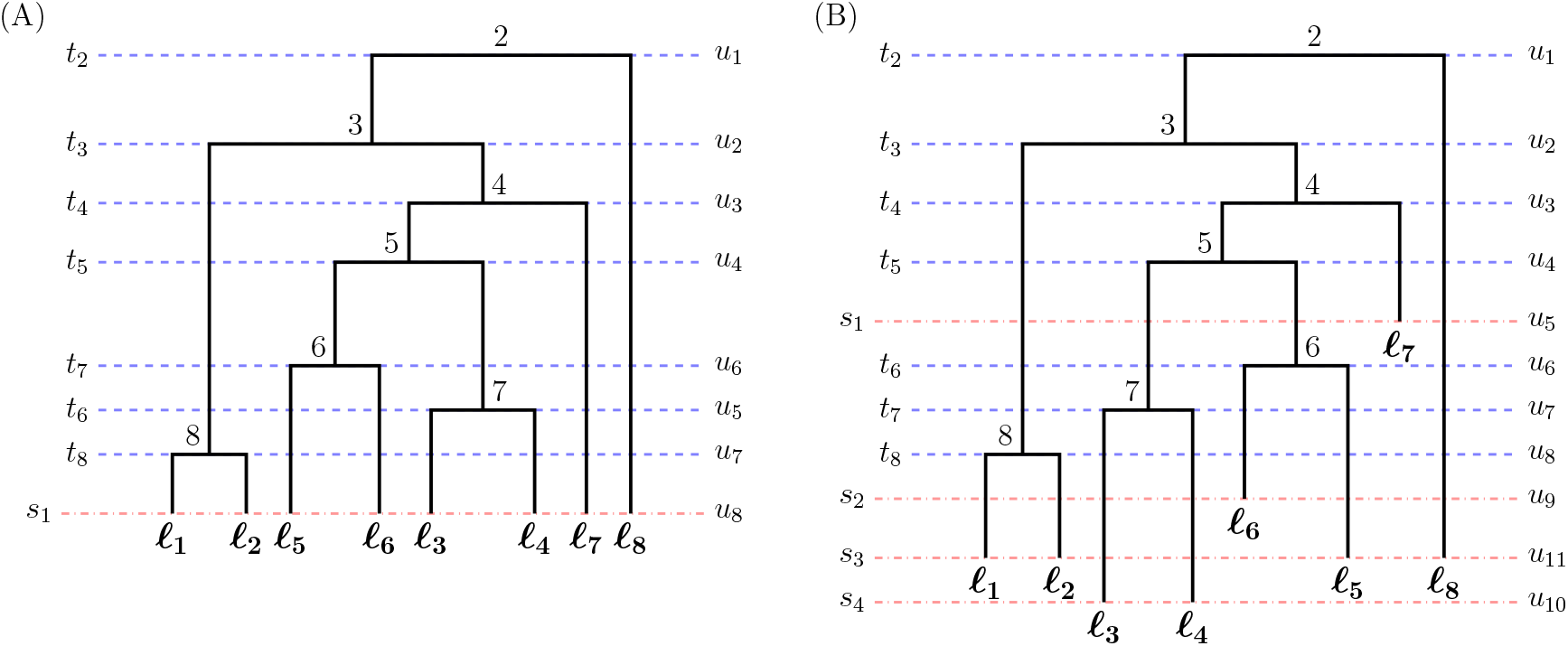
Unique labeling of ranked tree shapes and ranked genealogies. (A) Example of the unique labeling of a ranked genealogy with isochronous sampling. (B) Example of the unique labeling of a ranked genealogy with heterochronous sampling.

### Analysis of human influenza A virus from different continents

We selected all available sequences collected from January 2014 to January 2019 with complete HA segments in the GISAID EpiFlu database: 438, 410, and 510 sequences from New York, Chile, and Singapore, respectively. In order to compare trees with the same numbers of leaves, we kept 410 randomly selected sequences. We aligned the sequences using the FFT-NS-i option of MAFFT v7.427 software (Katoh et al., 2002; Katoh and Standley, 2013).

We used BEAST v1.10.4 (Suchard et al., 2018) to sample from the posterior distribution of variable effective population size trajectories and ranked and labeled genealogies with the following settings: SRD06 codon-partition substitution model (Shapiro et al., 2005) to accommodate rate heterogeneity among sites, strict molecular clock, and a Skygrid model with a regular grid of 100 points for inferring the effective population size trajectory (Gill et al., 2013). We ran the MCMC chain with 10^7^ steps and thinned every 10^4^.

### Software

- The source code for our distance metrics is available at https://github.com/JuliaPalacios/phylodyn.
- To perform MDS, we used the `cmdscale` function in R package stats.
- For our adaptation to the BHV metric, we used the Geodesic Treepath Problem (GTP) software implemented in Java (Owen and Provan, 2011).
- For the KC metric, we used `multiDist` implemented in the R package `treespace` (Kendall and Colijn, 2016).
- For the CP distance, we used the `vecMultiDistUnlab` in the R package `treetop` (Colijn and Plazzotta, 2018) with default parameters.
- To generated random isochronous ranked tree shapes, we used the `simulate_tree` function in the R package `apTreeshape` (Maliet et al., 2018) with the following parameters. In the beta-distribution simulation, *β ∈ {−*1.9, *−*1.5, *−*1, 0, 100}, *α* = 1, *ϵ* = 0.001, and *η* = 1. In the alpha-beta distribution simulation, *α ∈ {−*2, *−*1, 0, 1, 2}, *β* = 0, *ϵ* = 0.001 and *η* = 1.
- To generate random isochronous coalescent trees with different branch length distributions, we used the `rcoal` function from the `ape` package in R.
- To generate random heterochronous coalescent trees with different sampling events or different branch length distributions, we used the `coalsim` function implemented in `phylodyn` package in R.
- The preferential sampling of the hypothesized temperate region influenza dynamics was performed using the `pref_sample` function in phylodyn (Karcher et al., 2017).

## Acknowledgments

We would like to acknowledge Nina Miolane and Susan Holmes for useful discussion of distances and other embedding techniques. J.A.P. and N.A.R. acknowledge support from National Institutes of Health grant R01-GM-131404. J.A.P. acknowledges support from the Alfred P. Sloan Foundation.

## Supplementary Material

### S1 Alpha-Beta splitting model by Maliet et al. (2018)

**Algorithm 1.**
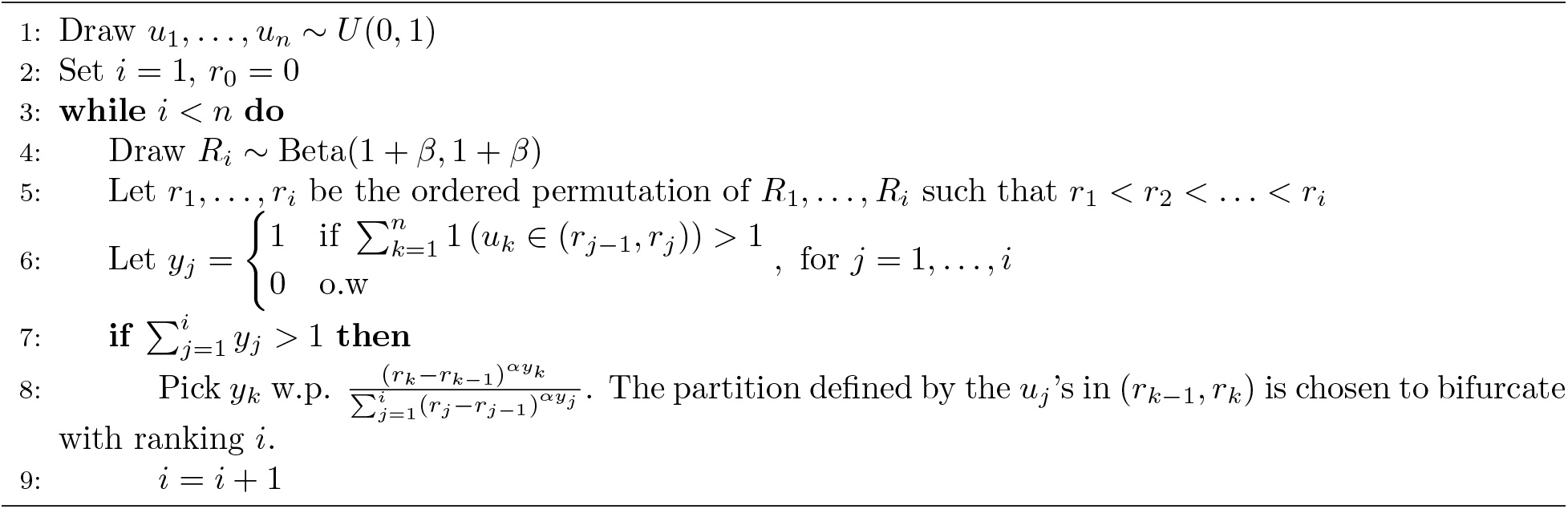
Simulation of a labeled ranked tree shape according to alpha-beta splitting model.

### S2 Proof of Theorem 1

*Proof*. Consider a ranked tree shape *T*^*R*^ with *n* leaves sampled at *m* different sampling times. We denote the total number of change points in *T*^*R*^ by *r* = *n*+*m*−1 and its ordered change point times by (*u_r_, u_r−_*_1_, *…, u*_1_), 0 = *u_r_* < *u_r−1_* < … < *u_1_*, with time increasing into the past. The internal nodes of *T*^*R*^ are labeled by the indices of their coalescent times, and all leaves of *T*^*R*^ are labeled by the indices of their sampling times. We note that for convenience, internal nodes are no longer labeled 2, …, *n* from the root to leaves, but they are labeled by their time-event indices (see Figure S1). Each internal node has a unique label, but the leaf nodes with the same sampling time share the same label. We define *N* = {1, …, *r*} to be a set of all node labels, *I* to be a set of all internal node labels, and *S* to be a set of all leaf node labels. Note that *I* and *S* are disjoint and contain *n* − 1 and *m* elements, respectively.

For *i ∈ I*, let *o_i_* = (*x*_*i,1*_, *x*_*i,2*_) denote the ordered pair of labels of the two immediate descendants of internal node *i*, such that *i < x_i,1_ ≤ x_i,2_*. We denote the set of all pairs *i* and *o_i_* = (*x_i,1_*, *x_i,2_*) choices in *T*^*R*^ by *X* = {(*i*, *o*_*i*_) | *i* ∈ *I*}. Then *X* completely specifies *T*^*R*^: *T*^*R*^ is a directed graph from the root to tips and *X* encodes its adjacency matrix and the order of the internal node indices *i ∈ I* determines internal node rankings.

We define a function *ϕ* : *X* → {0, 1, 2}^*r*−1^ as follows:

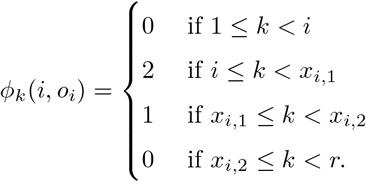

The *k*th element of *ϕ*(*i, o_i_*) is the number of immediate descendants of an internal node *i* present at the time interval (*u*_*k*+1_, *u_k_*). *ϕ* is an injective map. To prove this, let (*s, o_s_*), (*t, o_t_*) be two elements in *X* and let (*s, o_s_*) ≠ (*t, o_t_*). Because internal nodes of *T*^*R*^ are ranked, (*s, o_s_*) ≠ (*t, o_t_*) implies *s ≠ t*; without loss of generality, assume *s < t*. By the definition of the map *ϕ*, the *s*th element of *ϕ*(*s, o_s_*) is *ϕ_s_*(*s, o_s_*) = 2, while the *s*th element of *ϕ*(*t, o_t_*) is *ϕ_s_*(*t, o_t_*) = 0 because *s < t* and *t < x_t,1_ ≤ x_t,2_*. Thus, *ϕ*(*s, o_s_*) ≠ *ϕ*(*t, o_t_*) and *ϕ* is injective.

**Figure S1:**
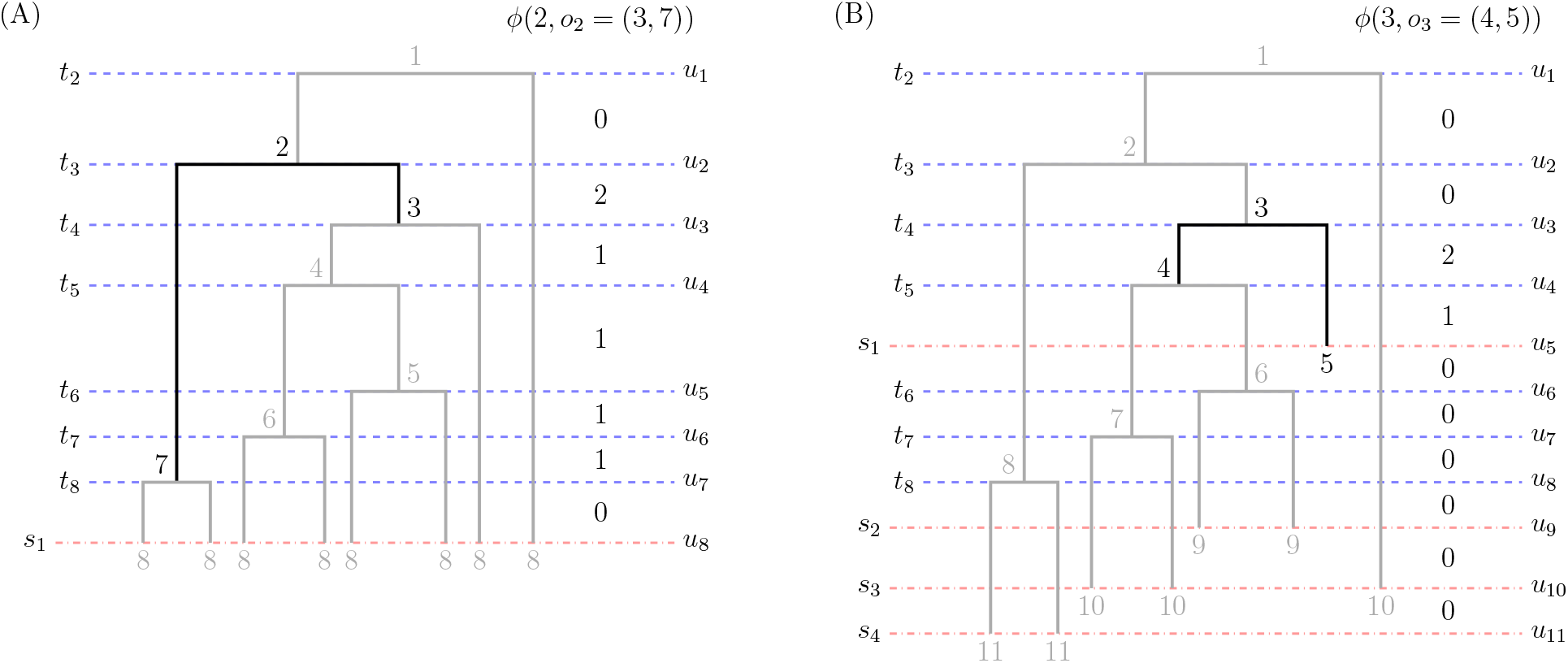
Example of *ϕ* mapping. (A) An example isochronous ranked tree shape. The set of internal node labels is *I* = 1, 2, 3, 4, 5, 6, 7 and the set of leaf node labels is *S* = 8 . For convenience, internal nodes are labeled by their time-event indices throughout the proof. The internal node with label 2 at time *u*_2_ has descendant nodes 3 and 7 at time *u*_3_ and *u*_7_, respectively (*o*_2_ = (3, 7)). The column vector *ϕ*(2, *o*_2_ = (3, 7)) = (0, 2, 1, 1, 1, 1, 0) indicates the number of direct descendants of node 2 at each change point time interval. (B) An example heterochronous ranked tree shape. The set of internal node labels is *I* = {1, 2, 3, 4, 6, 7, 8} and the set of leaf node labels is *S* = {5, 9, 10, 11}. The internal node with label 3 at time *u*_3_ has descendants node 4 and node 5 at time *u*_4_ and *u*_5_, respectively (*o*_3_ = (4, 5)). The column vector *ϕ*(3, *o*_3_ = (4, 5)) = (0, 0, 2, 1, 0, 0, 0, 0, 0, 0) indicates the number of direct descendants of node 3 at each change point time interval.

Let *η* : {1, *…, r − 1*} × {0, 1, 2}^*r*−1^ *→* {0, 1, 2}^*r*−1^ such that for **y** ∈ {0, 1, 2}^*r*−1^, the *j*-th element of *η* is

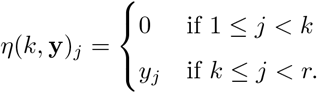

That is, *η*(*k*, **y**) sets all the first *k −* 1 entry values of **y** to 0. Note that the first *i −* 1 elements of *ϕ*(*i, o_i_*) are 0 by definition and thus, *η*(*i, ϕ*(*i, o_i_*)) = *ϕ*(*i, o_i_*).

Finally, for 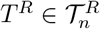, define 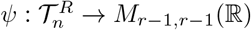, a function that maps a ranked tree shape with *n* leaves to a real valued square matrix of size *r −* 1, with *k*-th column:

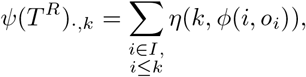

where *ψ*(*T*^*R*^)_*.,k*_ indicates the *k*th column of *ψ*(*T*^*R*^) and *I* is the set of all internal node labels as defined at the beginning of this section. By the definition of *η*, the first *k −* 1 values of the *k*th column *ψ*(*T*^*R*^)_*.,k*_ are 0, i.e., *ψ*(*T*^*R*^) is a lower triangular matrix. Because *ϕ* records the number of immediate descendants of a single internal node present at each time interval, *ψ*(*T*^*R*^)_*.,k*_ tracks the sum of all surviving immediate descendants of internal nodes with labels *i ≤ k* starting from time interval (*u_k_*_+1_, *u_k_*); thus, *ψ*(*T*^*R*^)_*s,k*_, with *k ≤ s*, represents the number of lineages of *T*^*R*^ in (*u_k_*_+1_, *u_k_*) that are still present at the time interval (*u_s_*_+1_, *u_s_*).

We prove that *ψ* is an injective map. Let 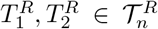 and 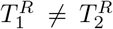. Because *X* = {(*i, o_i_*) | *i* ∈ *I*} completely specifies 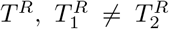 implies that there exists an index *ℓ ∈ {*1, *…, n −* 1} such that 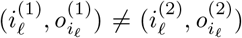. Here, *i*_ℓ_ indicates the *ℓ*th element of *I* sorted in increasing order. If there is more than one such index, choose *ℓ* to be the smallest of them. Without loss of generality, let 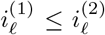. Then 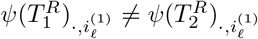 and thus 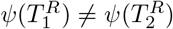.

Hence, *ψ* maps each ranked tree shape *T*^*R*^ to a unique matrix, i.e., given an **F**-matrix, if it encodes a ranked tree shape, it encodes exactly one ranked tree shape.

### S3 Proof of Proposition 2

*Proof.* The non-negativity and symmetry are trivial. The triangle inequality follows from the Minkowski inequality of *L*_1_ and *L*_2_ norms. It remains to prove the identity property: 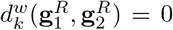 = 0 if and only if 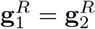 for *k* = 1, 2. It is clear that 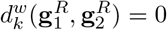 if 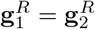 so we focus on 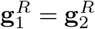 if 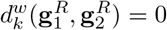.

The following proof is for 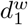. The proof for 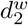 follows the same arguments.

We assume that the two genealogies have the same number of sampling events *m* and same number of leaves *n*, so that the **F**-matrices of 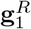 and 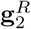 have the same dimension (*n* + *m −* 2) *×* (*n* + *m −* 2) dimension. We define *r* = *n* + *m −* 2 for notational simplicity.

Because we allow only a single event at each change time point *u_i_*, either coalescent or sampling, the first column of any **F**-matrix is (2, 1, *…,* 1) or (2, 1, *…,* 1, 0, *…,* 0). For the latter, we denote the row index of the last occurrence of 1 in the first column by 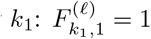 and 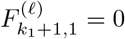 for some index 2 *≤ k*_1_ *≤ r* and ℓ = 1,2.

If **F**(1) and **F**(2) have different first columns, then for some index 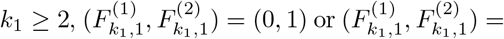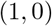. Because 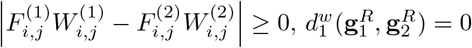 implies 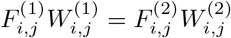 for all *i, j*. Therefore, 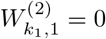 in the first case and 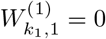 in the second case. However, this contradicts our assumption of positive time interval between two change points, and thus **F**^(1)^ and **F**^(2)^ must have the same First column.

If both **F**-matrices share the same first column, then 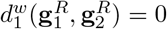 implies 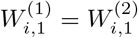 for all *i* = 1, …, r. Recalling *W*^*i,j*^ = *u*_*j*_ − *u*_*i*+1_, we have 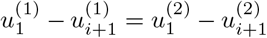. Because we assume 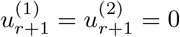, we can traverse through *i* in decreasing order starting from *i* = *r* to get 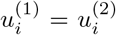 for all *i* = 1; … *r* + 1, which gives **W**^(1)^ = **W**^(2)^.

Along with **W**^(1)^ = **W**^(2)^, 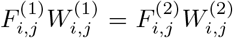 implies 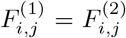 for all *i*, *j*, i.e **F**^(1)^ = **F**^(2)^, and thus 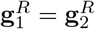

### S4 Embeddings in MDS

We chose MDS in two dimensions to visualize matrices of pairwise distances. In general, our metrics are well explained in the MDS visualization; however, the other distances are usually poorly represented in this space. In this section, we propose a measure of distortion and correlation to assess how well the embedding preserves the pairwise distances for each metric. In our examples, the distortion measure shown in Table S3 suggests that our *d*_2_ metric has the best MDS embedding in general of all distance functions considered with our *d*_1_ metric a close second. Similarly, the correlation measure shown in Table S4 confirms that our *d*_1_ and *d*_2_ metrics, with near perfect correlations, have far better embedding in the 2-dimensional MDS space than the other distances considered.

#### S4.1 Distortion

To assess our distances and its MDS embedding in two dimensions, we compute the following distortion statistic (Bourgain, 1985) defined as follows:

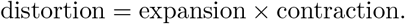

where expansion and contraction are defined as follows. For a given sample of ranked tree shapes 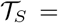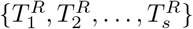 with *n* leaves in 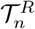,

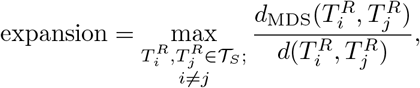

where *d*_MDS_ is the *L*_2_-Euclidean distance in the reduced MDS space and *d* is any distance function on ranked tree shapes, and

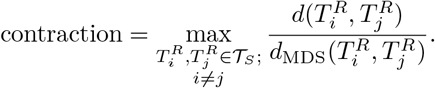

The distortion on the ranked genealogies is defined similarly. The comparison of distortions for simulated ranked tree shapes and ranked genealogies can be found in Table S3.

#### S4.2 Correlation

As a second measure for assessing our distances and its MDS embedding in two dimensions, we compute the Pearson correlation coefficient between the two vectors of pairwise distances between sampled ranked tree shapes, one from using any distance functions *d* on ranked tree shapes and the other from the *L*_2_-Euclidean distance *d*_MDS_ in the reduced MDS space:

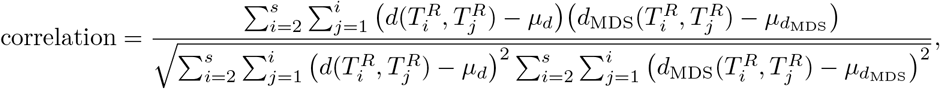

where *s* is the number of sampled ranked tree shapes. *µ_d_*MDS and *µ_d_* are the mean of the pairwise distances using *L*_2_-Euclidean distance in the MDS space and using any distance functions *d* on the sampled ranked tree shapes, respectively. The comparisons of correlations for simulated ranked tree shapes and ranked genealogies appear in Table S4.

## Supplementary Figures

**Figure S2:**
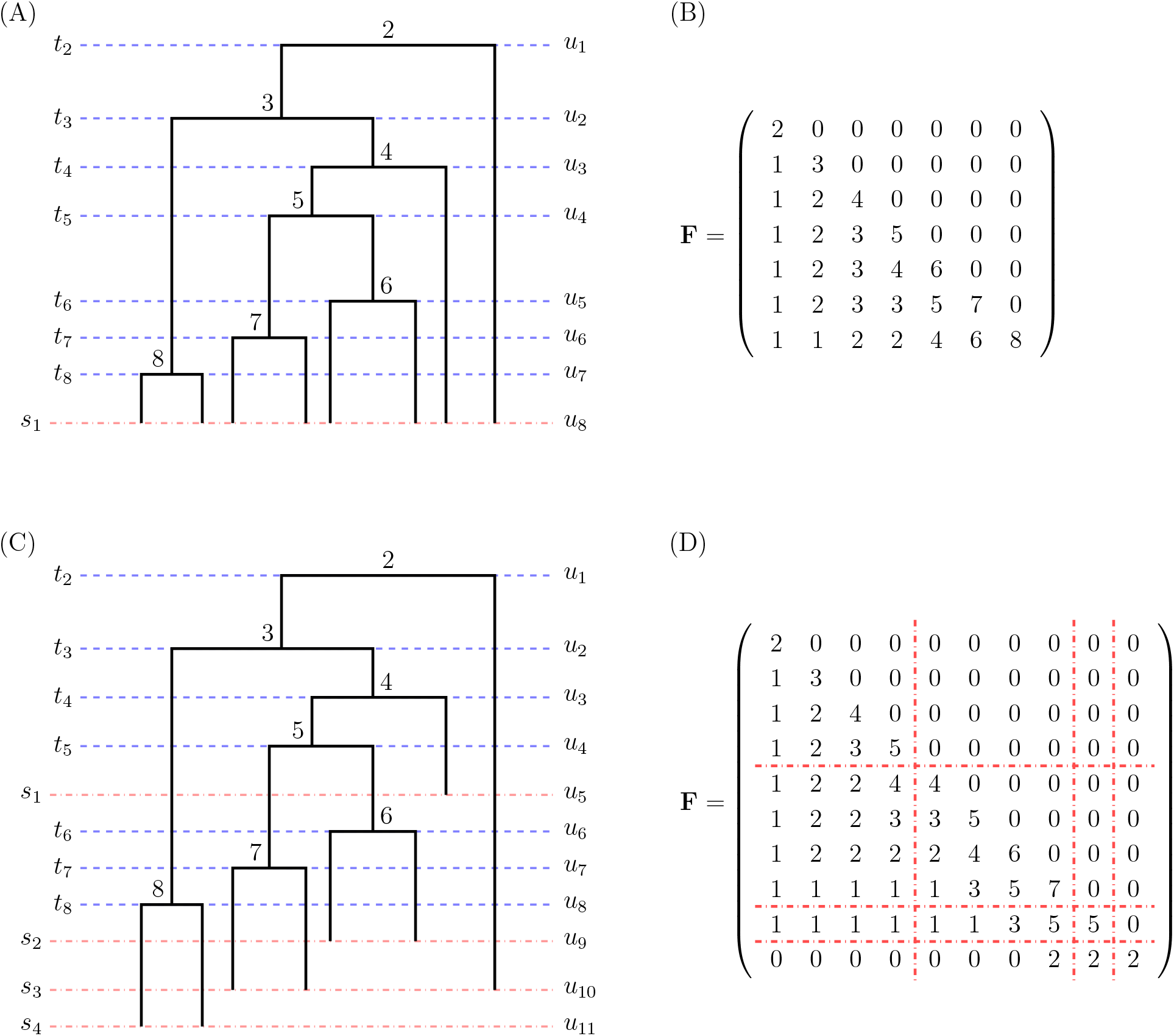
Bijection of ranked tree shapes and F-matrices for isochronous and heterochronous trees. (A) Example of a ranked genealogy with isochronous sampling. (B) The corresponding **F**-matrix that encodes the ranked tree shape information of the tree in (A). (C) Example of a ranked genealogy with heterochronous sampling. (D) The corresponding **F**-matrix of the heterochronous ranked tree shape in (C). Blue dotted lines indicate coalescent events and red dotted lines represent sampling events. In (C), coalescent times are denoted by 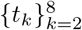, sampling times by 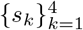, and the number of lineages changes at change points 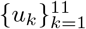.

**Figure S3:**
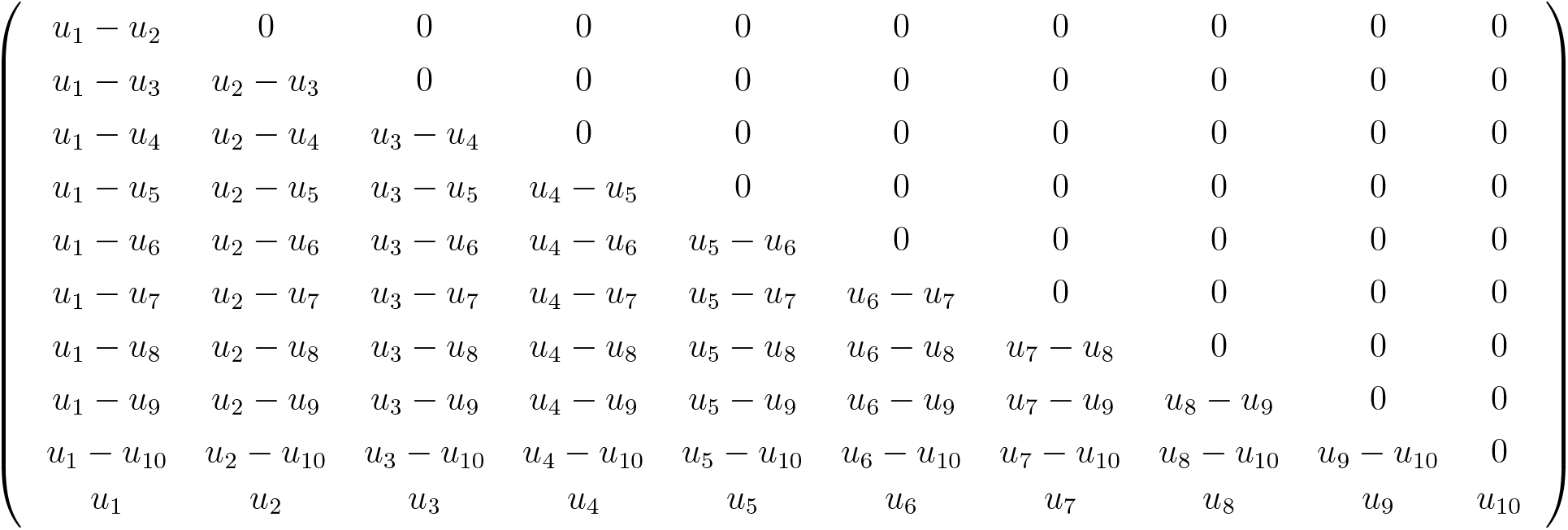
Example of the weight matrix W. The weight matrix associated with the example heterochronous ranked genealogy and its **F**-matrix in Figures S2(C) and (D). In the last row, *u*_11_ is suppressed because we set the initial sampling time to be *u*_11_ = 0.

**Figure S4:**
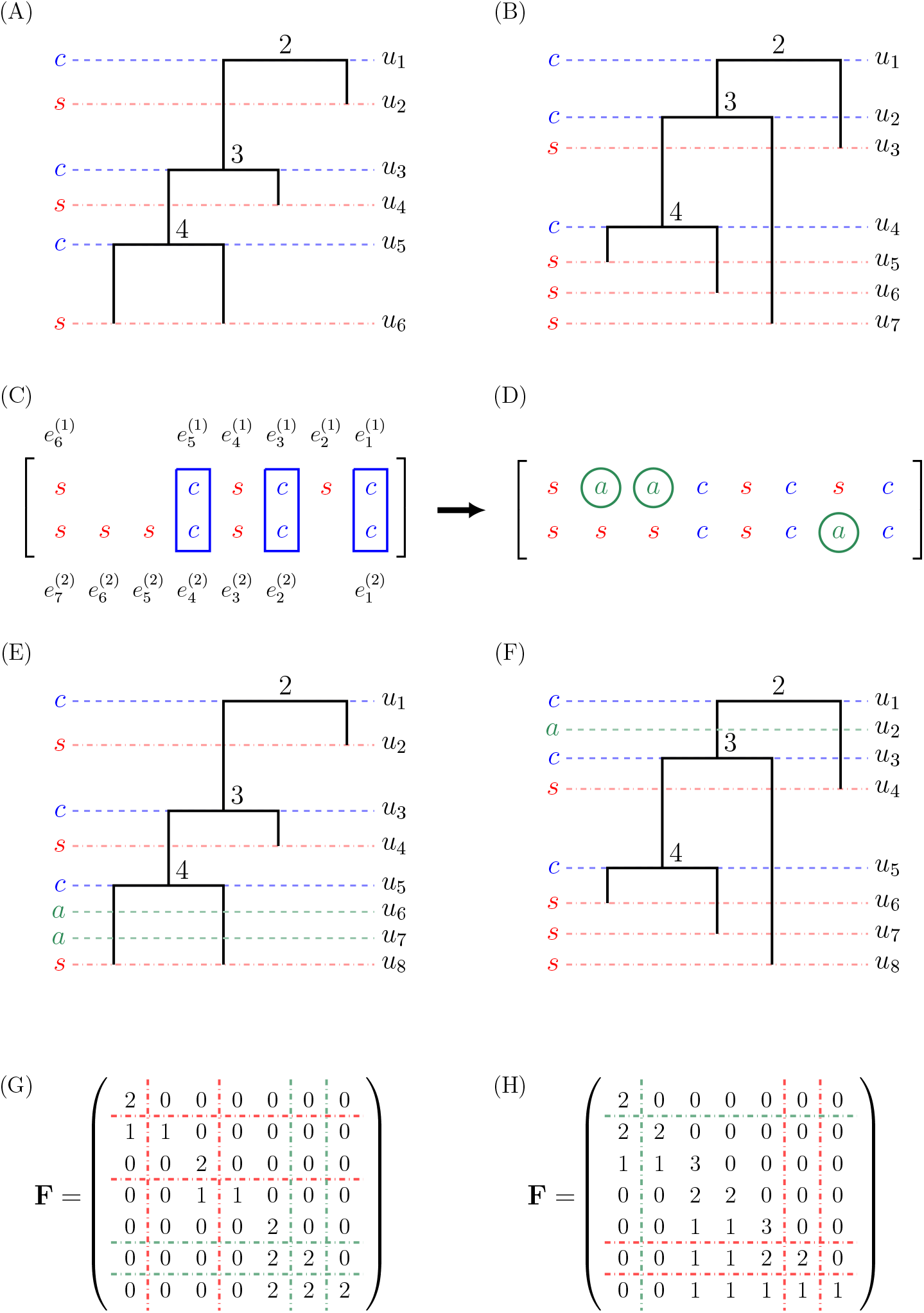
Example of augmented F-matrix representation of ranked tree shapes. In order to compute the *d*_1_ or *d*_2_ distances between ranked tree shapes with equally many samples but different numbers of sampling events, such as ranked tree shapes (A) and (B), we insert artificial sampling events with 0 samples in order to match their dimension. (A)-(B) Two ranked tree shapes of 4 samples with different sampling events. (C) Alignment of event vectors of two trees. The *n* − 1 coalescent events are aligned first by matching *i*th coalescent event of a tree to the *i*th coalescent event of the other tree (*i* = 1, …, *n* − 1). The sampling events are then matched by increasing index order in the event vector. (D) Augmentation of artificial sampling events *a* between coalescent events or between the first sampling and the first coalescent event. (E)-(F) Augmented ranked tree shapes. (G)-(H) Augmented **F**-matrix representations.

**Figure S5:**
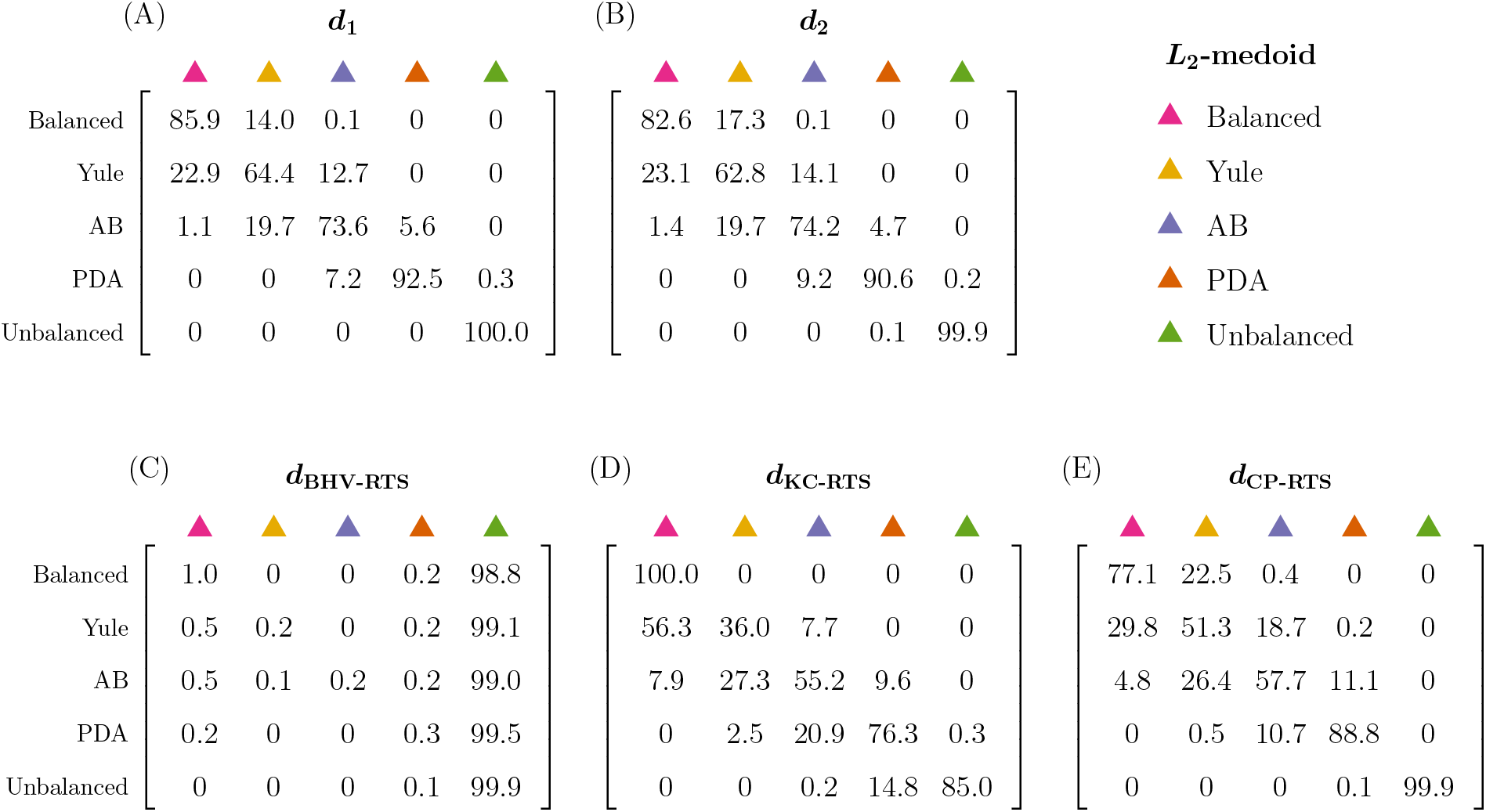
Comparison of metrics: discrimination of isochronous ranked tree shapes under different beta-splitting models. We compare the performance of different distances on ranked tree shapes according to how well they separate trees simulated from the beta-splitting distribution of ranked tree shapes with different balance parameters *β*. Rows indicate the sampling distribution and columns indicate the *L*_2_-medoid of each distribution. Each matrix corresponds to a different distance metric. Entry (*i, j*) corresponds to the percentage of trees simulated from the *i*-th distribution that are closer to the *j*-th *L*_2_-medoid than to the medoids of any other columns. The color scheme of the *L*_2_-medoids follows Figure 6. The mean diagonal values are 83.28, 82.02, 20.32, 70.50, and 74.96 for matrices (A)-(E), respectively.

**Figure S6:**
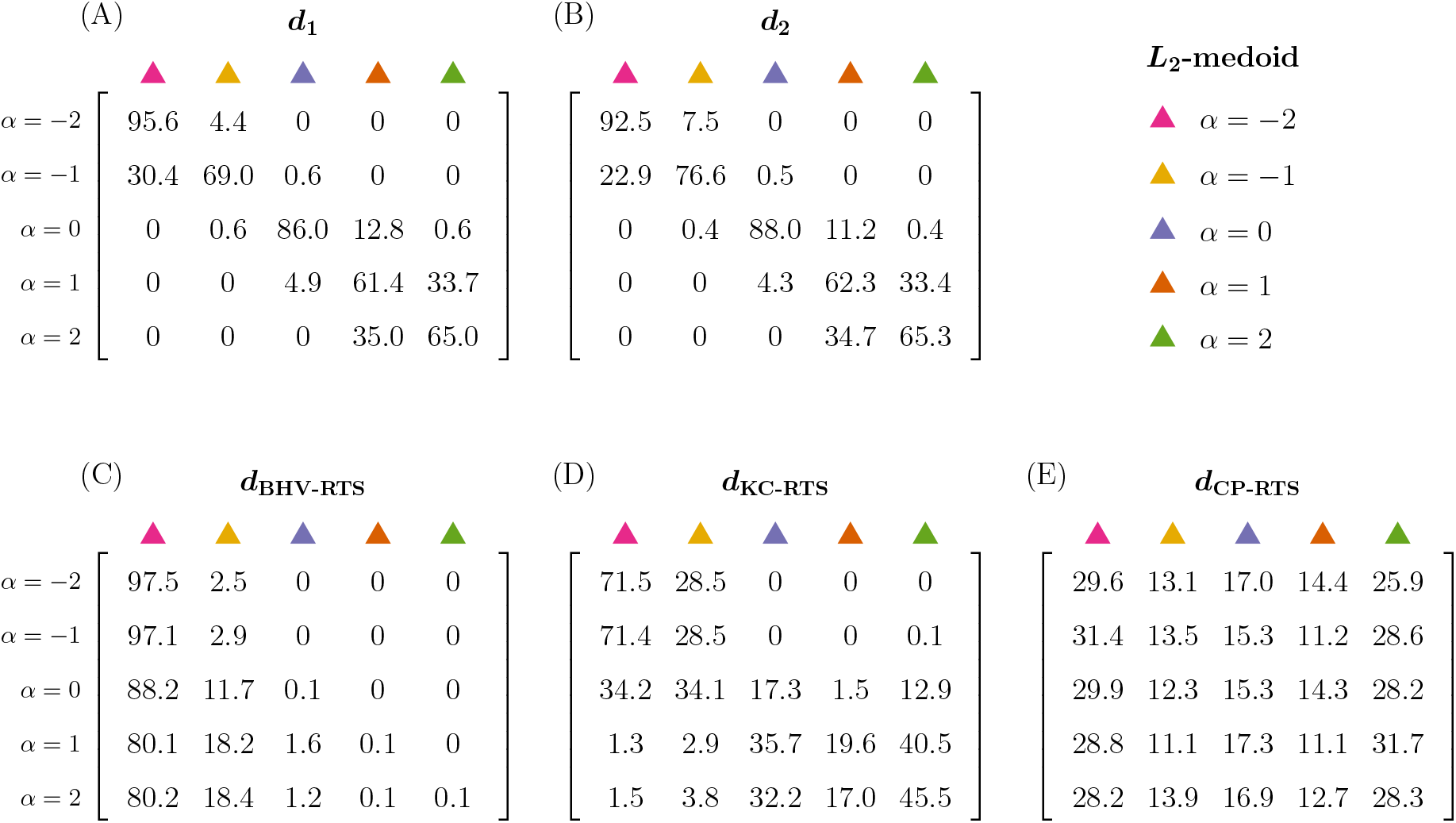
Comparison of metrics: discrimination of isochronous ranked tree shapes under different alpha-beta splitting models. We compare the performance of different distances on ranked tree shapes according to how well they separate trees simulated from the alpha-beta splitting distribution of ranked tree shapes with different parameter values *α* which regulates the internal node ranking of a given tree shape. The format of the matrices follows Figure S5. The simulation values and the color scheme of the *L*_2_-medoids follow Figure 7. The mean diagonal values are 75.40, 76.94, 20.14, 36.48, and 19.56 for matrices (A)-(E), respectively.

**Figure S7:**
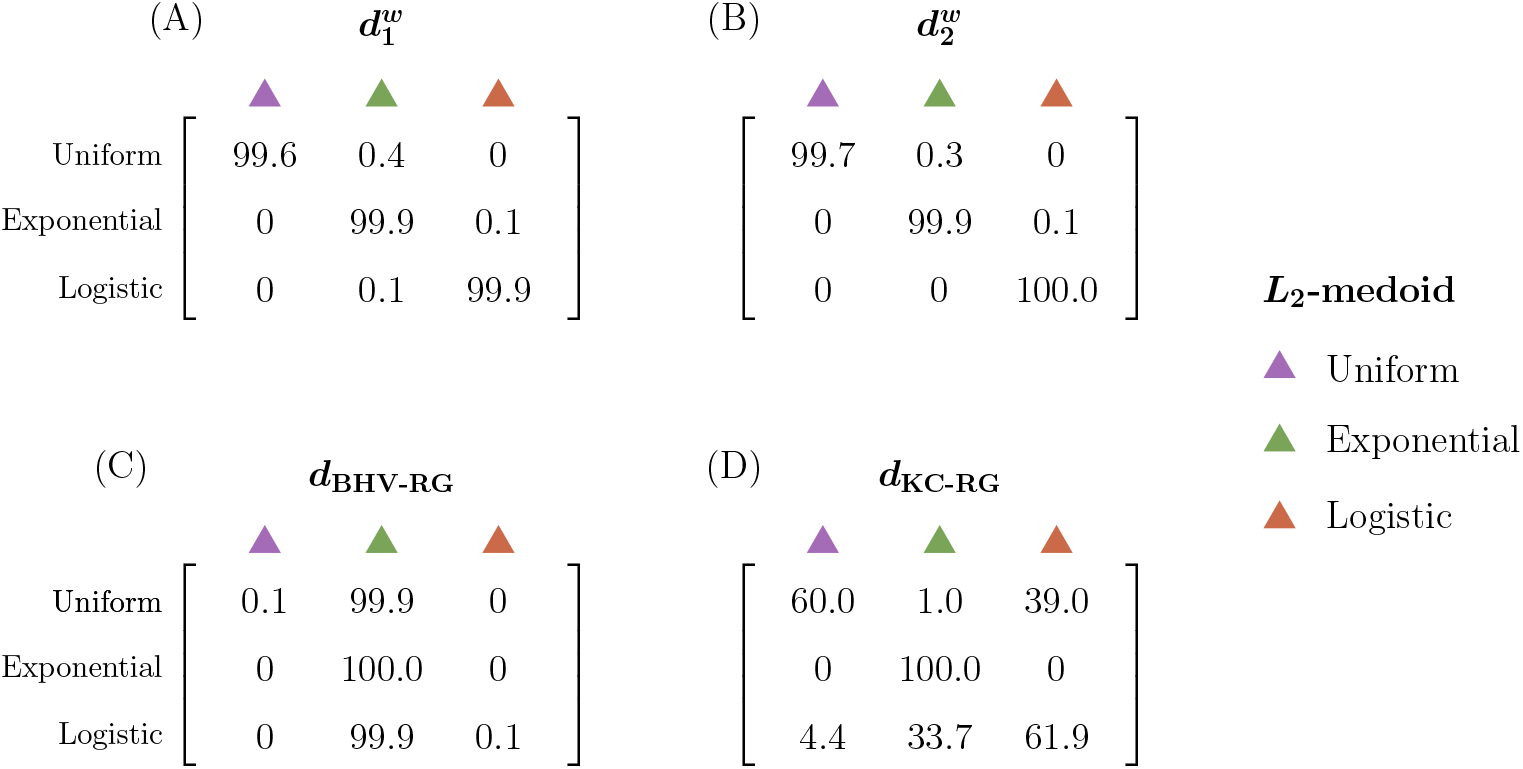
Comparison of metrics: isochronous ranked genealogies under different demographic models. We compare the performance of different distances on ranked genealogies according to how well they separate trees simulated from the *λ*(*t*)-coalescent with different population histories. The format of the matrices follows Figure S5. The simulation values and the color scheme of the *L*_2_-medoids follow Figure 8. The mean diagonal values are 99.80, 99.87, 33.40, and 73.97 for matrices (A)-(D), respectively.

**Figure S8:**
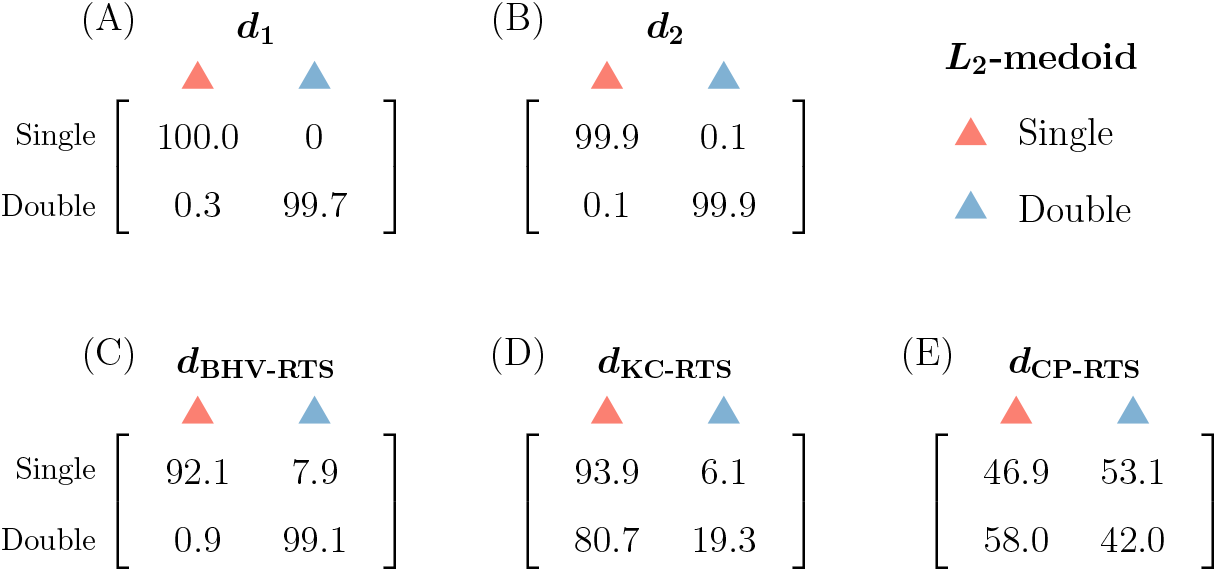
Comparison of metrics: heterochronous ranked tree shapes with different sampling schemes. We compare the performance of different distances on ranked genealogies according to how well they separate trees simulated from the heterochronous Tajima coalescent with different sampling sequences **s** and **n**. The format of the matrices follows Figure S5. The simulation values and the color scheme of the *L*_2_-medoids follow Figure 9. The mean diagonal values are 99.85, 99.90, 95.60, 56.60, and 44.45 for matrices (A)-(E), respectively.

**Figure S9:**
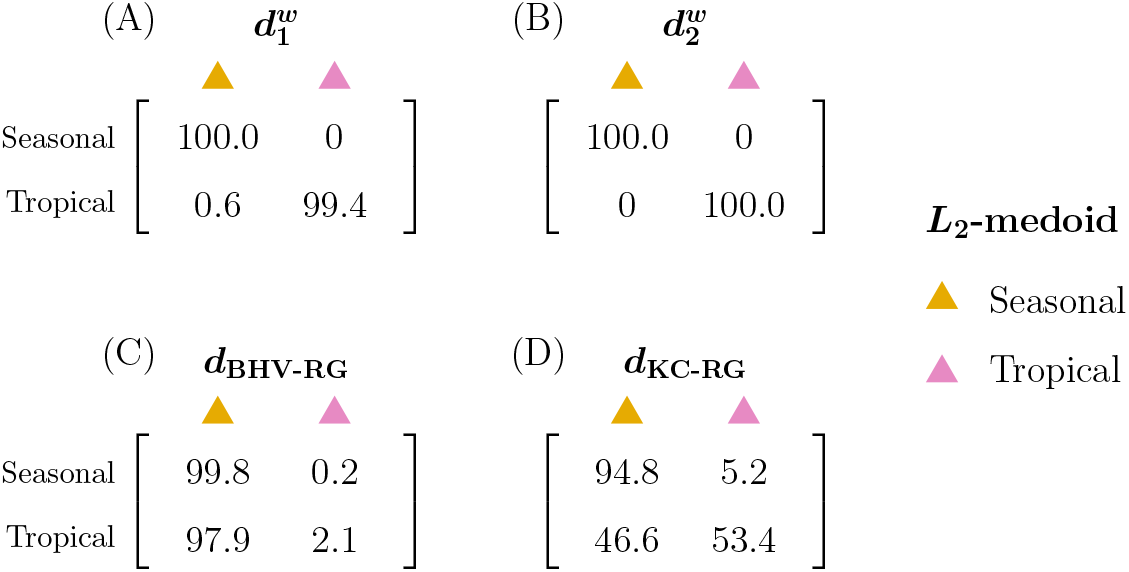
Comparison of metrics: heterochronous ranked genealogies with different simulation models. We compare the performance of different distances on ranked genealogies according to how well they separate trees simulated from the heterochronous *λ*(*t*)-coalescent with different population histories and sampling sequences **s** and **n**. The format of the matrices follows Figure S5. The simulation values and the color scheme of the *L*_2_-medoids follow Figure 10. The mean diagonal values are 99.70, 100.00, 50.95, and 74.10 for matrices (A)-(D), respectively.

**Figure S10:**
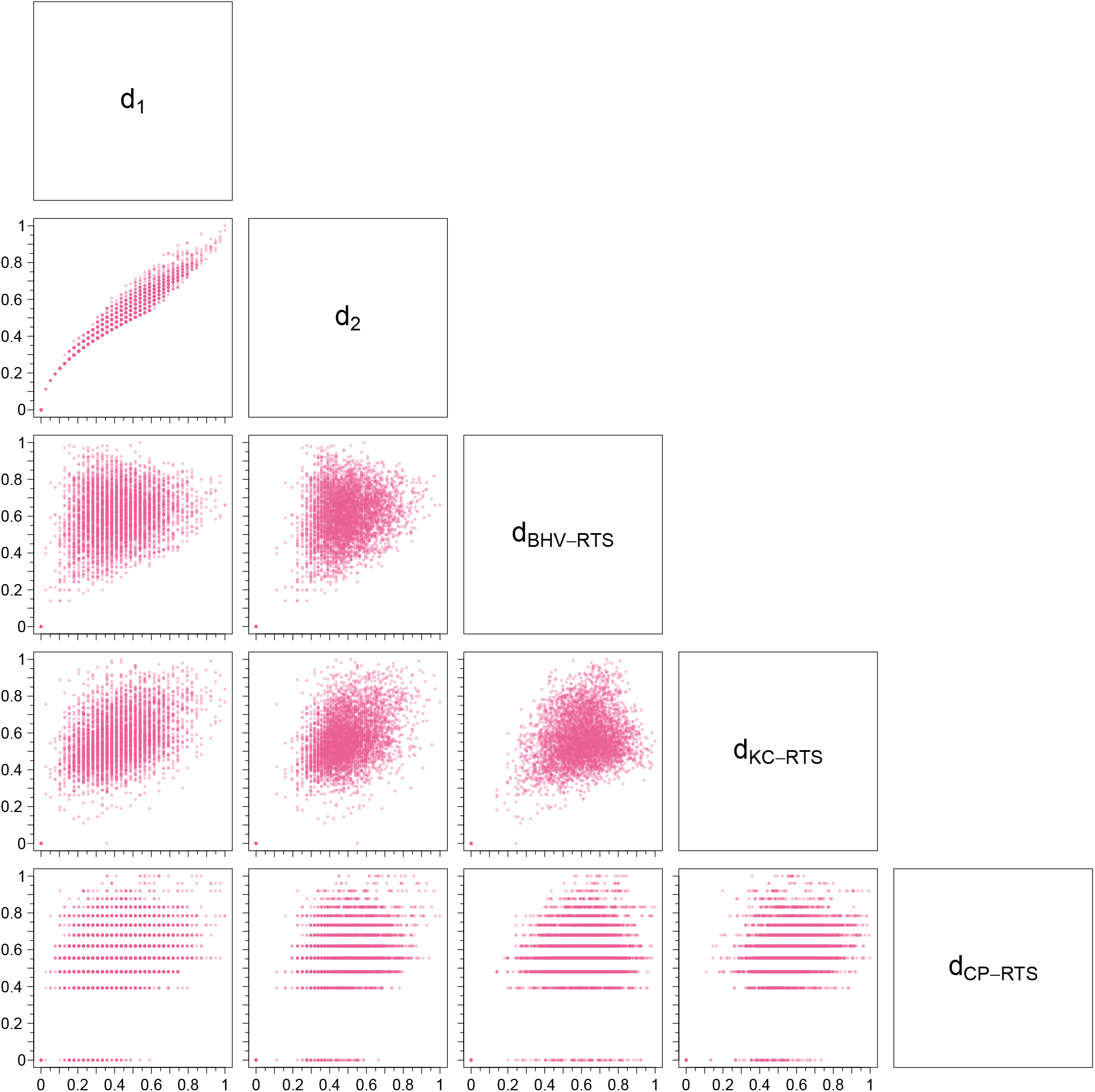
Pairwise comparisons of metrics using 100 simulated ranked tree shapes with *n* = 10 leaves. Each plot contains 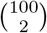 points, where each point represents two pairwise distances between simulated ranked tree shapes. *d*_CP-RTS_ displays the lowest resolution among all the metrics plotted. Multiple pairs of different ranked tree shapes share the same *d*_CP-RTS_ value while many of those pairs have distinct values using the other metrics.

**Figure S11:**
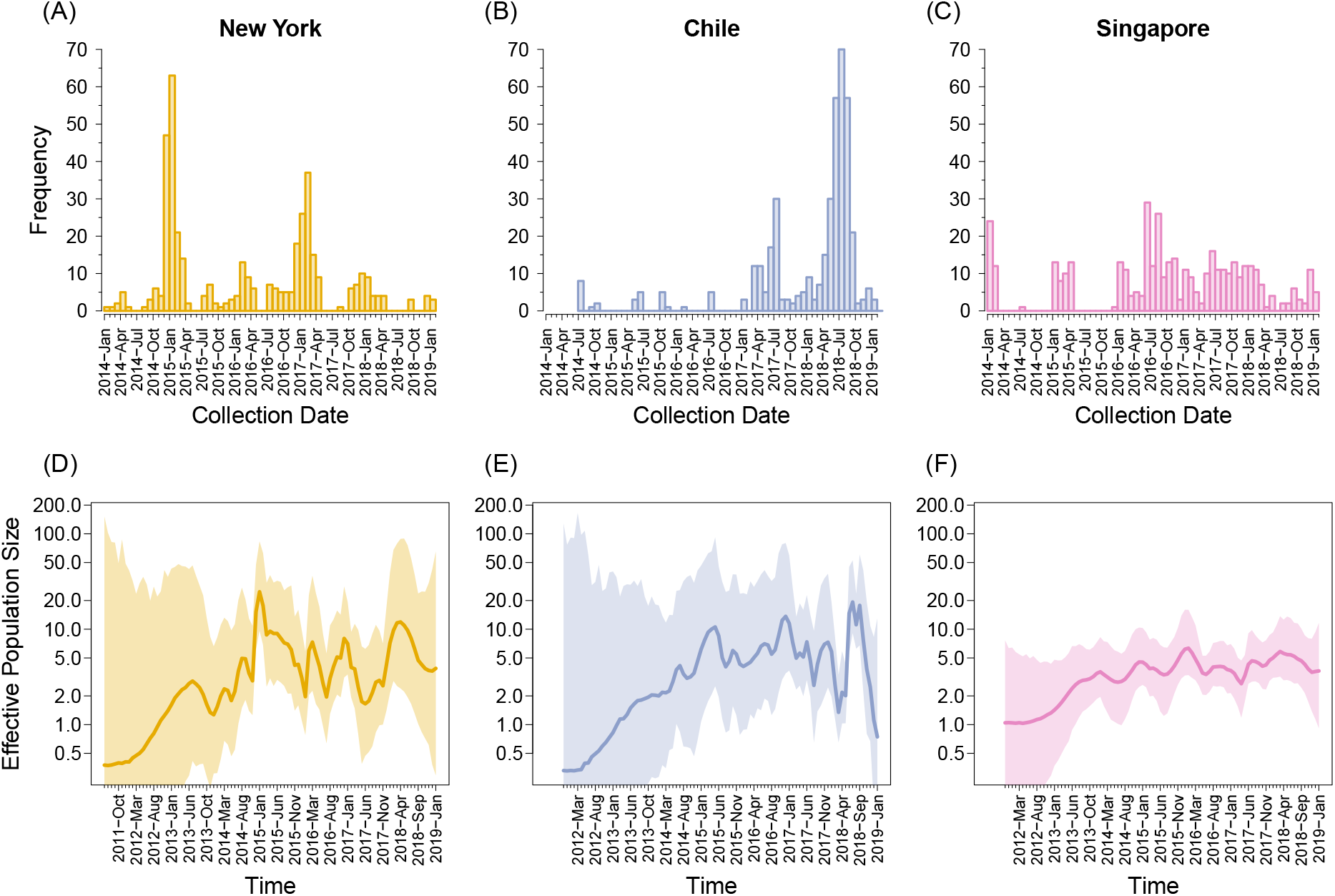
Human influenza A virus collection dates and inferred effective population size trajectories. (A) Collection date histogram, New York; (B) Collection date histogram, Chile; (C) Collection date histogram, Singapore; (D) Inferred population size trajectory with BEAST, New York; (E) Chile; and (F) Singapore. Data description and color scheme follow Figure 12.

## Supplementary Tables

**Table S1.** Summary of dispersion comparisons for ranked tree shapes. Comparison of dispersion of ranked tree shape distribution (Section S4.1) using distance functions between ranked tree shapes. (A) Isochronous ranked tree shapes simulated from the beta-splitting model with varying *β* parameters (Figure 6). (B) Isochronous ranked tree shapes simulated from the alpha-beta splitting model with varying *α* parameters and fixed *β* = 0 (Figure 7). (C) Heterochronous ranked tree shapes simulated from different sampling schemes (Figure 9).

| | $d_1$ | $d_2$ | $d_{\text{BHV-RTS}}$ | $d_{\text{KC-RTS}}$ | $d_{\text{CP-RTS}}$ |
| --- | --- | --- | --- | --- | --- |
| Balanced | 5541.21 | 117.14 | 377.84 | 128.20 | 8.51 |
| Yule | 5889.31 | 119.40 | 363.33 | 182.52 | 8.97 |
| AB | 6579.90 | 127.37 | 345.82 | 271.78 | 9.58 |
| PDA | 6966.50 | 133.24 | 304.49 | 490.98 | 10.48 |
| Unbalanced | 4391.36 | 90.88 | 154.44 | 944.90 | 12.50 |

(B) Isochronous ranked tree shapes, alpha-beta splitting model
| | $d_1$ | $d_2$ | $d_{\text{BHV-RTS}}$ | $d_{\text{KC-RTS}}$ | $d_{\text{CP-RTS}}$ |
| --- | --- | --- | --- | --- | --- |
| $\alpha = -2$ | 4701.27 | 90.81 | 85.33 | 110.76 | 8.95 |
| $\alpha = -1$ | 7202.89 | 135.70 | 110.64 | 126.06 | 9.00 |
| $\alpha = 0$ | 7719.71 | 150.57 | 262.18 | 172.73 | 8.96 |
| $\alpha = 1$ | 6084.73 | 122.77 | 363.17 | 182.49 | 9.06 |
| $\alpha = 2$ | 5668.18 | 115.80 | 376.98 | 179.18 | 9.08 |

(C) Heterochronous ranked tree shapes
| | $d_1$ | $d_2$ | $d_{\text{BHV-RTS}}$ | $d_{\text{KC-RTS}}$ | $d_{\text{CP-RTS}}$ |
| --- | --- | --- | --- | --- | --- |
| Single | 10359.43 | 277.88 | 350.26 | 230.17 | 9.47 |
| Double | 10423.86 | 238.90 | 324.47 | 257.65 | 9.62 |

**Table S2.** Summary of dispersion comparisons for ranked genealogies. Comparison of dispersion of ranked genealogies using distance functions between ranked genealogies. (A) Isochronous ranked genealogies simulated from different population trajectories under neutral coalescent model (Figure 8). (B) Heterochronous ranked genealogies simulated from different population trajectories under neutral coalescent model Figure 10). (C) Heterochronous ranked genealogies of human influenza A virus data from different continental groups (Figure 12).

| | $d_1$ | $d_2$ | $d_{\text{BHV-RTS}}$ | $d_{\text{KC-RTS}}$ |
| --- | --- | --- | --- | --- |
| Constant | 6420771.00 | 142726.04 | 33007.42 | 391150.52 |
| Exponential | 1331068.00 | 24847.09 | 1397.17 | 4743.48 |
| Logistic | 1744504.00 | 39291.83 | 8751.45 | 109258.59 |

(B) Heterochronous ranked genealogies
| | $d_1$ | $d_2$ | $d_{\text{BHV-RTS}}$ | $d_{\text{KC-RTS}}$ |
| --- | --- | --- | --- | --- |
| Seasonal | 98912.28 | 1074.80 | 105.49 | 1335.42 |
| Tropical | 307851.92 | 2703.92 | 344.64 | 4363.61 |

(C) Heterochronous ranked genealogies, human influenza A virus
| | $d_1$ | $d_2$ | $d_1^w$ | $d_2^w$ |
| --- | --- | --- | --- | --- |
| New York | 668449.00 | 3982.78 | 139378.00 | 501.75 |
| Chile | 947326.20 | 4896.46 | 181856.90 | 568.68 |
| Singapore | 373360.50 | 1775.27 | 172247.40 | 544.05 |

**Table S3.** Summary of distortion comparisons. Comparison of distortion (Section S4.1) using distance functions between ranked trees shapes and ranked genealogies.

| | $d_1$ | $d_2$ | $d_{\text{BHV-RTS}}$ | $d_{\text{KC-RTS}}$ | $d_{\text{CP-RTS}}$ |
| --- | --- | --- | --- | --- | --- |
| Isochronous<br>ranked tree shapes<br>(beta splitting) | 5582.02 | 2369.98 | 8586.00 | 14331.64 | 27721.77 |
| Isochronous<br>ranked tree shapes<br>(alpha-beta splitting) | 3979.43 | 3028.30 | 10941.54 | 23989.64 | 5446.20 |
| Heterochronous<br>ranked tree shapes | 2343.93 | 2969.33 | 4553.04 | 12034.03 | 1247.20 |

| | $d_1^w$ | $d_2^w$ | $d_{\text{BHV-RG}}$ | $d_{\text{KC-RG}}$ |
| --- | --- | --- | --- | --- |
| Isochronous<br>ranked genealogies | 6751.44 | 1836.89 | 18569.28 | 13716.09 |
| Heterochronous<br>ranked genealogies | 2283.54 | 2503.3 | 24424.35 | 3214.8 |

**Table S4.** Summary of correlations. Comparison of correlation (Section S4.2) between original distances and Euclidean distances in 2-dimensional MDS comparisons.

| | $d_1$ | $d_2$ | $d_{\text{BHV-RTS}}$ | $d_{\text{KC-RTS}}$ | $d_{\text{CP-RTS}}$ |
| --- | --- | --- | --- | --- | --- |
| Isochronous<br>ranked tree shapes<br>(beta splitting) | 0.998 | 0.998 | 0.401 | 0.982 | 0.988 |
| Isochronous<br>ranked tree shapes<br>(alpha-beta splitting) | 0.998 | 0.999 | 0.648 | 0.518 | 0.909 |
| Heterochronous<br>ranked tree shapes | 0.984 | 0.986 | 0.634 | 0.734 | 0.898 |

| | $d_1^w$ | $d_2^w$ | $d_{\text{BHV-RG}}$ | $d_{\text{KC-RG}}$ |
| --- | --- | --- | --- | --- |
| Isochronous<br>ranked genealogies | 0.981 | 0.954 | 0.527 | 0.946 |
| Heterochronous<br>ranked genealogies | 0.926 | 0.888 | 0.377 | 0.924 |

